# The role of ER exit sites in maintaining P-body organization and transmitting ER stress response during *Drosophila melanogaster* oogenesis

**DOI:** 10.1101/2024.07.03.601952

**Authors:** Samantha N. Milano, Livia V. Bayer, Julie J. Ko, Caroline E. Casella, Diana P. Bratu

**Affiliations:** Department of Biological Sciences, Hunter College, City University of New York, NY, 10065 USA; Program in Molecular, Cellular, and Developmental Biology, The Graduate Center, City University of New York, NY, 10016 USA

## Abstract

Processing bodies (P-bodies) are cytoplasmic membrane-less organelles which host multiple mRNA processing events. While the fundamental principles of P-body organization are beginning to be elucidated *in vitro*, a nuanced understanding of how their assembly is regulated *in vivo* remains elusive. Here, we investigate the potential link between ER exit sites and P-bodies in *Drosophila melanogaster* egg chambers. Employing a combination of live and super-resolution imaging, we found that P-bodies associated with ER exit sites are larger and less mobile than cytoplasmic P-bodies, indicating that they constitute a distinct class of P-bodies which are more mature than their cytoplasmic counterparts. Moreover, we demonstrate that altering the composition of ER exit sites has differential effects on core P-body proteins (Me31B, Cup, and Trailer Hitch) suggesting a potential role for ER exit sites in P-body organization. We further show that in the absence of ER exit sites, P-body integrity is compromised and the stability and translational repression efficiency of the maternal mRNA, *oskar*, are reduced. Finally, we show that ER stress is communicated to P-bodies via ER exit sites, highlighting the pivotal role of ER exit sites as a bridge between membrane-bound and membrane-less organelles in ER stress response. Together, our data unveils the significance of ER exit sites not only in governing P-body organization, but also in facilitating inter-organellar communication during stress, potentially bearing implications for a variety of disease pathologies.

## INTRODUCTION

Investigations of biological condensates are unlocking insights into a broad array of fields ranging from mRNA regulation and stress response to long-distance transcript localization. While the breadth of condensate function is beginning to be elucidated, little is known about how condensate assembly is regulated *in vivo*. *In vitro* studies have revealed the ability of mRNA and proteins to form condensates *de novo*, via Liquid Liquid Phase Separation (LLPS) (1–4). Such condensation events are driven by weak mRNA:mRNA, mRNA:protein, and protein:protein interactions (5,6). Many proteins capable of LLPS have intrinsically disordered regions that allow for multiplexing weak interactions (7,8). Under physiological conditions, these weak interactions generate a local concentration of mRNAs and proteins which is significantly higher than the surrounding cytoplasm. Due to this disparity, it becomes thermodynamically favorable for a granule to form a distinct liquid phase (5,9). Once formed, these liquid-like granules can mature and become solid-like (10,11). Whether this bottom-up mechanism is the only driving factor that dictates condensate formation in complex tissues is yet to be deciphered. Here, we look to highlight the influence of specialized regions of the ER, ER exit sites (ERES), on condensate formation in the *Drosophila melanogaster* female germline.

The *D. melanogaster* ovary represents a complex tissue consisting of 16 to 20 chains of egg chambers. The germline of each egg chamber is a 16-cell cyst that shares a common cytoplasm. Early in development, one of these cells becomes the oocyte, and the other 15 subsequently become nurse cells. The oocyte remains transcriptionally silent while the nurse cells produce all the transcripts required for the oocyte’s development (12). These transcripts must be stably maintained and translationally repressed as they are spatio-temporally localized to the distant oocyte (13). One method by which this silenced transport is achieved is via processing bodies (P-bodies). P-bodies play active roles in the mRNA life cycle, such as hosting transcript storage, translational repression, and mRNA decay (14,15).

P-bodies are differentiated from similar cytoplasmic granules, i.e. stress granules, by their constitutive presence in the cell’s cytoplasm (15). The number of P-bodies varies depending on cell type, further hinting at their functional importance. Oocytes and neurons both contain a prodigious number of P-bodies. As these are polarized cells where transport of stable and translationally repressed transcripts occurs over long distances, it suggests that P-bodies may play an important role in this transport process (16,17). Interestingly, these cells also utilize localized ERES along the ER membrane.

ERES are specialized regions of the ER where COPII vesicles form, allowing for protein secretion (18). Recent evidence has indicated that these sites are not always stochastically located along the ER membrane; in some cell types, their distribution can be spatio-temporally regulated (19). Increased ERES markers have been observed preferentially at the end of axons (20). Similarly, in the *D. melanogaster* egg chamber, it has been shown that protein localization is regulated in part via selection of localized ERES (21).

The functional link between the ER and P-bodies is not novel (22). Previous studies in *D. melanogaster* egg chambers have demonstrated that core P-body proteins, Me31B (RNA helicase), Cup (eIF4E -BP), and Trailer Hitch (Tral) (LSM 14-like) colocalize with the ER membrane (23,24). These studies further revealed that Tral directly binds the mRNAs of two COPII proteins: Sar1 and Sec13, and in the absence of Tral protein, ERES are aberrantly localized indicating that P-bodies play a role in ERES regulation during *D. melanogaster* oogenesis (23). In yeast, the homolog of Tral, Car-1, similarly governs proper ER organization, implying that this may be a conserved mechanism of spatio-temporal protein regulation via targeted secretion sites (25). As evidence continues to mount for the spatio-temporal regulation of ERES by P-bodies, recent reports have also surfaced indicating the reciprocal regulation of P-bodies by the ER (26).

ER contact sites with membrane-bound organelles have been noted (27). Now, the possibility that the ER also forms contact sites with membrane-less organelles is emerging. In yeast, P-body components are directly associated with the ER membrane, and blocking ER secretion leads to an increase in P-body formation at the ER (28). In cell culture, the ER has been shown to directly facilitate P-body fission, further bolstering the idea that the interface between the ER and P-bodies has meaningful biological consequences (29).

Here, we look to further delineate the relationship between ERES and P-body assembly *in vivo*. Using super-resolution imaging and RNAi-facilitated knockdowns, we found that P-bodies at ERES are distinct from cytoplasmic P-bodies. Furthermore, upon knockdown of COPII components, we observed alterations in key P-body protein condensates, suggesting a potential involvement of COPII proteins in overall condensate organization. Moreover, in the absence of ERES, P-bodies did not form detectable condensates. Finally, our studies unveiled that ERES stress may be communicated via P-bodies rather than stress granules, a finding that could have implications for diseases ranging from cancer to neurodegeneration (30,31).

## RESULTS

### P-bodies colocalized with ER exit sites are distinct from cytoplasmic P-bodies

P-bodies are heterogeneous in composition, size, and morphology (32). Since P-bodies have been shown to interface with the ER (28), we sought to quantitatively characterize the distinctions between these ER-associated P-bodies and cytoplasmic P-bodies. We assessed two common condensate parameters: sphericity and volume. Based on thermodynamics, liquid-like droplets with small volumes are spherical due to the influence of surface tension. Over time, these condensates can coalesce, with a tendency to increase in volume to reduce interfacial tension. Liquid-like condensates can also mature as stronger intra-condensate interactions develop making them more gel-like and less spherical (33,34).

We chose to visualize P-bodies marked with endogenous Me31B-GFP and the ER by expressing UAS-KDEL-RFP induced with the Gal4/UAS system (Fig. 1A). These flies were viable over multiple generations, suggesting that there are no catastrophic phenotypes associated with the transgene. Interestingly, we found that Me31B-GFP condensates associated with the ER were, on average, 80% larger (0.973*µ*m^3^ compared to 0.196*µ*m^3^) and 12% less spherical than their cytoplasmic counterparts (Fig. 1B and 1C). This result is consistent with these granules being more mature as they are larger and more structured. Next, we sought to determine the specific region of the ER that was responsible for the difference in condensate characteristics.

**Figure 1:**
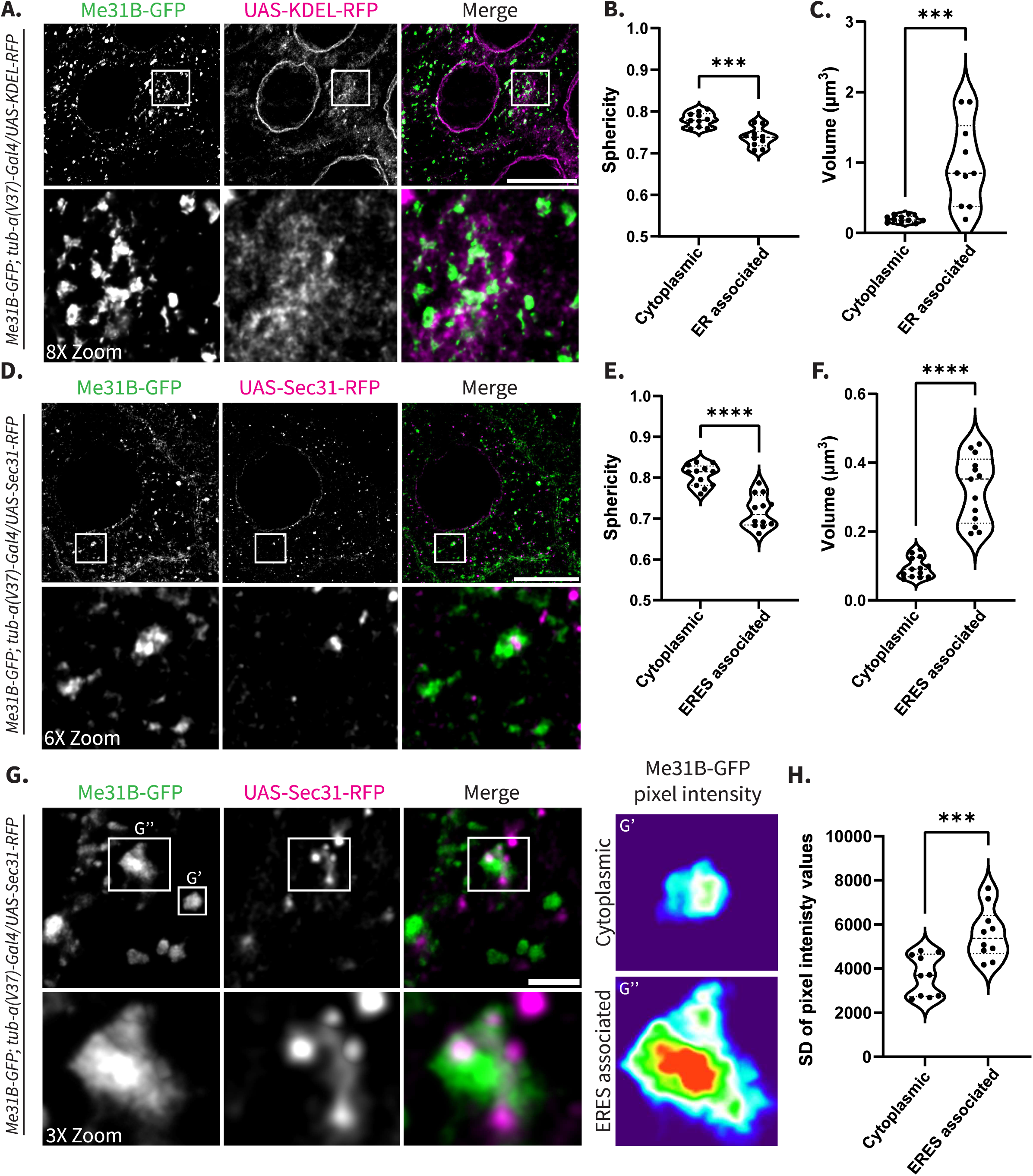
P-bodies colocalized with ER exit sites are distinct from cytoplasmic P-bodies. **(A)** Endogenous Me31B-GFP colocalized with UAS-KDEL-RFP in nurse cell cytoplasm. Images are XY projections of 5 optical Z slices of 0.3*µ*m. Scale bars are 20*µ*m and 2.5*µ*m respectively in the zoomed inset. **(B,C)** Sphericity and volume measurements comparing cytoplasmic and ER-associated P-bodies (n=10). **(D)** Endogenous Me31B-GFP colocalization with UAS-Sec31-RFP in nurse cell cytoplasm. Images are XY projections of 5 optical Z slices of 0.3*µ*m. Scale bars are 20*µ*m and 3.3*µ*m respectively in the zoomed inset. **(E,F)** Sphericity and volume measurements of cytoplasmic and ERES-associated P-bodies (n=10). **(G)** Visualization Me31B-GFP with UAS-Sec31-RFP via STED. Images are XY projections of 5 optical Z slices of 0.22*µ*m. Scale bars are 2*µ*m and 670nm respectively in the zoomed inset. **(G’,G’’)** Intensity heat map of a cytoplasmic and ERES-associated P-body respectively. **(H)** Standard deviation of pixel intensity values for cytoplasmic and ERES-associated P-bodies (n=10). For all plots, each data point represents the average value of all P-bodies detected in an image. Significance was assessed using a Mann-Whitney statistical test. **** p< .0001.

We chose to focus on ERES-associated P-bodies because of the previous connection between P-bodies and ERES (23,25,26). We conducted the same analysis using Me31B-GFP and a Gal4-induced ERES protein, Sec31-RFP (Fig. 1D). To ensure that the transgene was an accurate read-out of ERES localization, we used a COPII antibody to verify that Sec31-RFP localizes properly (Fig. S1). We found that ERES-associated P-bodies were on average 71% larger (0.324*µ*m^3^ compared to 0.095*µ*m^3^) and 25% less spherical than cytoplasmic P-bodies, further suggesting that these P-bodies represent a distinct class of P-bodies (Fig. 1E and 1F).

As the liquid-like state of condensates has been shown to influence P-body function (35), we asked if ERES-associated P-bodies exhibit a different physical state than cytoplasmic P-bodies and therefore may perform different functions. To address this question, we employed STED super-resolution microscopy to assess ‘surface texture’ as an indicator of condensate physical state (Fig. 1G). ‘Rougher’ P-bodies indicate more solid/gel-like condensates as fluorescence signal is distributed in clumped structures, while ‘smoother’ P-bodies indicate more liquid-like condensates, with more evenly dispersed fluorescence signal.

To quantify fluorescence distribution, we utilized standard deviation (SD) of pixel intensity (36). This technique requires assessing the intensity of each pixel within the boundary of a P-body and obtaining a SD which indicates the range of pixel intensities within the condensate. Since ‘rough’ P-bodies contain fluorescence peaks and valleys, with high and low pixel intensities respectively (Fig. 1G’’), they reflect a larger SD of pixel intensity when compared to ‘smooth’ P-bodies which present pixels with more homogenic intensity values (Fig. 1G’). For each detected P-body (above 0.05*µ*m^3^), a SD of pixel intensity values was calculated.

Using this analysis, we found that ERES-associated P-bodies were 34% ‘rougher’ than cytoplasmic P-bodies (Fig. 1H). As condensates can mature and ‘harden’ over time, this may indicate that these P-bodies are older and forming gel-like condensate states and therefore may perform a different function (10,11).

### ERES-associated P-bodies are dynamically distinct from cytoplasmic P-bodies

P-bodies display multiple patterns of dynamics; some stay stationary, others move within local vicinities, while some move along the microtubule network (37). To attain a better understanding of how ERES-associated P-bodies differ from cytoplasmic P-bodies, we performed live imaging to visualize Me31B-GFP with Sec31-RFP. All P-bodies not associated with Sec31-RFP were binned as cytoplasmic P-bodies (Fig. 2A), while all P-bodies with Sec31-RFP were binned as ERES-associated P-bodies (Fig. 2B). Interestingly, ERES-associated P-bodies did not appear to move long distances and were often imaged throughout the entirety of the live imaging session, while cytoplasmic P-bodies were only imaged over a few time points, indicating that they move long distances. We calculated track linearity and found that cytoplasmic P-bodies have 46% more linear tracks compared to ERES-associated P-bodies (Fig. 2C). This data indicates that ERES-associated P-bodies move continuously in a local area while cytoplasmic P-bodies feature directed movement. Based on these results, we hypothesized that cytoplasmic P-bodies are moving via the microtubule network while ERES-associated P-bodies exhibit Brownian motion.

**Figure 2:**
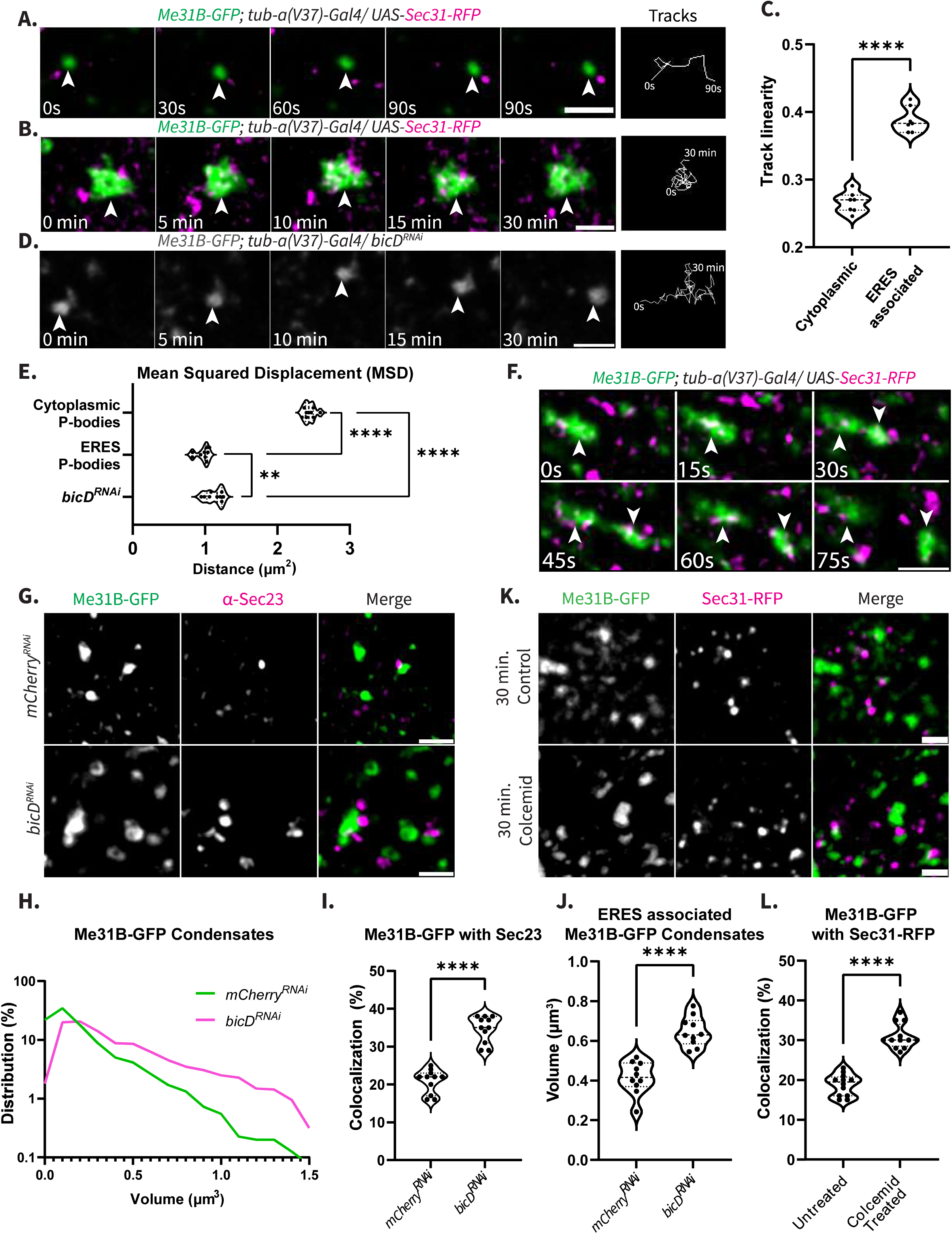
ERES-associated P-bodies are dynamically distinct from cytoplasmic P-bodies. **(A)** Tracking of a Me31B-GFP-labeled P-body not colocalizing with Sec31-RFP. **(B)** Tracking of Me31B-GFP-labeled P-body colocalizing with Sec31-RFP. **(C)** Track linearity calculated for cytoplasmic and ERES-associated P-bodies (n=7). **(D)** Tracking of Me31B-GFP in *bicD^RNAi^* nurse cell. Images acquired every 15 sec for 30 min. **(E)** Mean Squared Displacement of cytoplasmic P-bodies, ERES-associated P-bodies, and P-bodies in a *bicD^RNAi^* background over a period of 45 sec (n=9). **(F)** P-body fission event at ERES. Arrows represent separate bodies. **(G)** Me31B-GFP and Sec23 co-visualized with immunofluorescence in *mCherry^RNAi^* and *bicD^RNAi^* knockdown backgrounds. XY projections of 5 optical Z slices of 0.3*µ*m. **(H)** Me31B-GFP condensate volume distribution in *mCherry^RNAi^* and *bicD^RNAi^* backgrounds. X-axis is shown on a log scale. **(I)** Volume of Me31B-GFP condensates colocalized with Sec23 in *mCherry^RNAi^* control and *bicD^RNAi^* egg chambers (n=11). **(J)** Colocalization analysis of Me31B-GFP with Sec23, calculated with “Surface Detection” on Imaris software (n=10). **(K)** Me31B-GFP with Sec31-RFP incubated in Schneider’s media control or 10*µ*M colcemid for 30 minutes. XY projections of 5 optical Z slices of 0.3*µ*m. **(L)** Colocalization analysis of Me31B-GFP with Sec31-RFP (n=10). All scale bars are 2*µ*m. For all plots, each point represents the average value of all P-bodies in an image. Significance was calculated using a Mann-Whitney statistical test. **** p< .0001.

Our hypothesis is supported by a recent study that identified Me31B as part of the Egalitarian protein interactome, suggesting that P-bodies move along microtubules via the Egalitarian/BicD/Dynein complex (38). To determine if the cytoplasmic P-bodies’ movements relied on association with microtubules, we knocked down BicD using RNAi and assessed Me31B-GFP particle dynamics (Fig. 2D). The knockdown efficiency was confirmed via immunofluorescence (Fig. S2A). Notably, in the *bicD^RNAi^* background, Me31B-GFP condensates exhibited movement patterns resembling those of ERES-associated P-bodies and remained in frame for extended durations. Calculations of track mean squared displacement (MSD) revealed that cytoplasmic P-bodies traveled 2.5 times further than ERES-associated P-bodies within a 45 sec timeframe. Furthermore, P-bodies in the *bicD^RNAi^* background displayed MSD more comparable to ERES-associated P-bodies than cytoplasmic P-bodies, suggesting that cytoplasmic P-bodies do utilize the microtubule network (Fig. 2E).

Interestingly, we observed that ERES-associated P-bodies underwent instances of fission (Fig. 2F). As thermodynamics dictate that condensates should increase in size, this fission is likely due to an outside force. This led us to postulate that cytoplasmic P-bodies may be pulled away from ERES-associated P-bodies by the microtubule network. To investigate this, we visualized Me31B-GFP and Sec31-RFP in a *bicD^RNAi^* background (Fig. 2G). Microtubule expression was unaffected in the *bicD^RNAi^* egg chambers so alterations in cytoplasmic crowding did not affect our results (Fig. S2B). Interestingly, upon BicD knockdown, there was a noticeable shift in P-body distribution towards larger volumes (Fig. 2H) and Me31B-GFP colocalization with ERES increased by 66% (Fig. 2I). Together this indicates that, without connection to microtubules, the interaction between P-bodies and ERES is more stable, suggesting that perhaps the microtubule network is responsible for pulling P-bodies away from ERES. To buttress this finding, we assessed the volume of ERES-associated Me31B-GFP condensates in the *bicD^RNAi^* background and observed that they were 55% larger than in *mCherry^RNAi^* control egg chambers (0.646*µ*m^3^ compared to 0.416*µ*m^3^), suggesting that they may be undergoing fewer fission events (Fig. 2J).

Moreover, in egg chambers where microtubules were destabilized with 10*µ*M colcemid for 30 min, condensate distribution was similarly altered (Fig. S2C). ERES-associated P-bodies represented a larger percentage of the condensate population, 65% more than in the non-treated control (Fig. 2K and 2L), further suggesting that microtubules play a role in either forming or maintaining the cytoplasmic population of P-bodies. Taken together this data suggests that ER-associated P-bodies are a distinct class of P-bodies which we call ERES P-bodies.

### COPII vesicle proteins affect the organization of putative P-body protein condensates

As ERES P-bodies were quantitatively distinct from cytoplasmic P-bodies, we sought to better decipher the connection between P-bodies and ERES components. ERES are composed of a suite of conserved proteins. Sar1, a cytoplasmic GTPase, is thought to govern the assembly of ERES (39). Upon interaction with the membrane-bound GEF, Sec12, the GTP-bound Sar1 inserts into the membrane and facilitates deformation. By inducing curvature, Sar1 insertion enhances the affinity of additional Sar1 and propagates vesicle formation (40). In *D. melanogaster,* Sec16, a large scaffolding protein, is recruited to ERES independently of Sar1 (41). Once these two proteins are at the ER membrane, they assemble the inner COPII vesicle coat made up of Sec23 and Sec24 which subsequently recruits the outer-coat of Sec13 and Sec31 (Fig. 3A) (18).

**Figure 3:**
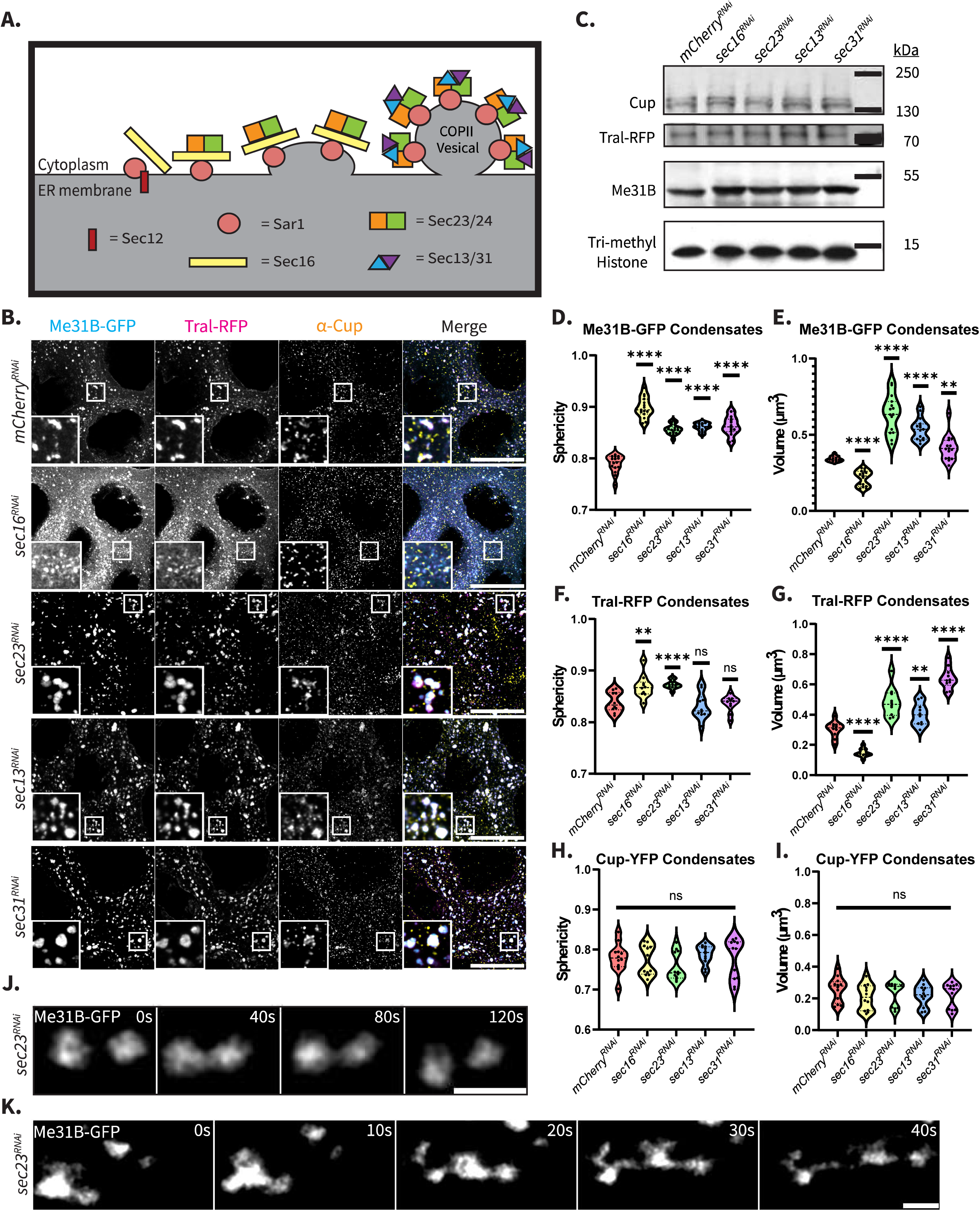
COPII vesicle proteins affect the organization of putative P-body protein condensates. **(A)** Schematic of ERES formation. **(B)** Colocalization of Me31B-GFP, Tral-RFP, and Cup in egg chambers with each COPII component knocked down. XY projections of 5 optical Z slices of 0.3*µ*m. Scale bars are 20*µ*m. **(C)** Western blot of Cup, Tral-RFP, and Me31B. Tri-methyl-Histone -- loading control. **(D,E)** Sphericity and volume measurements of Me31B-GFP condensates (n = 15). **(F,G)** Sphericity and volume measurements of Tral-RFP condensates (n = 10). **(H,I)** Sphericity and volume measurements for Cup-YFP condensates (n= 12). All calculations were performed on slides with only one fluorescently tagged protein. Significance was calculated using a Mann-Whitney statistical test. **** p< .0001. **(J,K)** Live imaging of Me31B-GFP in *sec23^RNAi^* egg chambers. Scale bars are 2*µ*m.

To probe the relationship between P-bodies and ERES, we separately analyzed three endogenously tagged P-body proteins: Me31B, Tral, and Cup, in the background of all publicly available COPII protein RNAi lines: *sec16^RNAi^*, *sec23^RNAi^*, *sec13^RNAi^*, and *sec31^RNAi^* (Fig. 3B). The RNAi line efficiencies were validated with antibodies when available, or RT-qPCR (Fig. S3A-D). Through this analysis, we discerned distinct roles for ERES proteins in modulating both the volume and morphology of P-body protein condensates, where the overall levels of P-body proteins were unaffected (Fig. 3C).

In *sec16^RNAi^* egg chambers, Me31B-GFP condensates were 37% smaller and 28% more spherical compared to condensates in *mCherry^RNAi^* egg chambers (Fig. 3D and 3E), indicating that Me31B-GFP condensates are either unable to fully form, or maintain, proper condensate state (35,42). Despite many P-bodies being smaller than those in control knockdowns, there was a population of larger condensates. We believe this is due to the incomplete efficiency of the *sec16^RNAi^*knockdown, although further experiments would help determine the cause of this bifurcated population (Fig. S3A). Similarly, when we analyzed Tral-RFP condensates in this background, they were on average 51% smaller and 7% more spherical than in control egg chambers (Fig. 3F and 3G). This suggests that Me31B and Tral may require efficient COPII vesicle formation to properly assemble into mature condensates.

We proceeded to knock down the inner-coat protein, Sec23, which is also the GTPase-activating protein (GAP) responsible for terminating COPII budding (43). In *sec23^RNAi^*egg chambers, Me31B-GFP condensates were 86% larger than in the control background (Fig. 3D). These condensates were also 17% more spherical (Fig. 3E). Similarly, Tral condensates were 59% larger and 8% more spherical compared to condensates in the control. This indicates that P-body maintenance is severely compromised, and overall P-body organization is aberrant in the *sec23^RNAi^* background.

Lastly, we investigated the outer-coat proteins: Sec13 and Sec31. The P-bodies in these knockdown backgrounds closely phenocopied the Sec23 knockdown, as Me31B-GFP condensates were 58% larger in the *sec13^RNAi^* egg chambers and 21% larger in *sec31^RNAi\^* egg chambers compared to control (Fig. 3D). Me31B-GFP condensates in these backgrounds were also on average 18% and 19% more spherical respectively (Fig. 3E). Similarly, Tral-RFP condensates were on average 33% and 122% larger than in the control egg chambers, however, did not display sphericities that differed significantly (Fig. 3F and 3G).

The formation of large condensates in Sec23, Sec13, and Sec31 knockdown egg chambers may arise from the role of these ERES proteins in terminating COPII vesicle formation. The interaction between Sec23 and Sar1 triggers the detachment of Sar1 from the membrane, allowing COPII vesicles to bud off. This activity is facilitated partly by Sec13 stabilization and more directly by Sec31’s role in activating Sec23’s GAP function (44,45).

Interestingly, neither the volume nor sphericity of Cup-YFP condensates were significantly altered in any of the COPII knockdown backgrounds (Fig. 3H and 3I). This finding was unexpected as it suggests that Cup may enter P-bodies via a distinct mechanism from Me31B and Tral, despite all three being known to form a ribonucleoprotein (RNP) complex (46,47). This indicates that ERES may play a crucial role in dictating P-body organization.

To ascertain the physical state of the large condensates observed in the coat protein knockdowns, we visualized Me31B-GFP in a *sec23^RNAi^* background, as this combination presented the most pronounced phenotype. Interestingly, we observed condensates moving towards each other, transiently coalescing, and subsequently separating. This suggests that these condensates may not exhibit purely liquid-like behavior, as they did not fuse upon contact (Fig. 3J). Notably, we visualized instances of condensates undergoing fission, potentially indicating that these condensates possess solid/gel-like cores surrounded by liquid-like shells, enabling them to separate even if they cannot completely fuse (Fig. 3K).

Given the previously demonstrated influence of the ER on microtubule arrangement, which in turn may affect P-body formation, we postulated that the disparity in condensate organization may arise from aberrant microtubule arrangement in the ERES knockdown backgrounds (48,49). To test for such pleiotropic effects, we visualized the integrity of microtubules, actin, and nuclear membranes in *sec16^RNAi^, sec23^RNAi^, sec13^RNAi^,* and *sec31^RNAi^* backgrounds and found that they were unaffected, suggesting that the condensate alterations were not a result of cellular breakdown (Fig. S3F). We also considered that the phenotypes may result from gross alterations in the ER. However, visualization of KDEL-RFP indicated that the overall ER membrane was unaffected in the knockdowns (Fig. S3G).

Finally, we considered that the presence of large condensates might be a consequence of stress. This is plausible as stress granules contain many of the same proteins as P-bodies. To assess this, we visualized the distribution of Rasputin (Rin), a stress granule marker, in each of the COPII protein knockdown backgrounds. Notably, we did not observe any Rin-GFP condensates, indicating the absence of stress granule formation (Fig. S3H). Taken together, these findings lead us to believe that condensate mis-organization is likely a direct consequence of improper COPII assembly.

### In the absence of ER exit sites, P-body integrity is compromised

Since our data suggests that COPII vesicle components play a crucial role in overall P-body organization, we aimed to visualize P-bodies in a background without ERES formation initiation. This was achieved by knocking down Sar1, verified with RT-qPCR (Fig. S4A). To confirm the absence of ERES formation, we visualized Sec23 and observed a substantial reduction in COPII puncta (Fig. S4B).

Since egg chambers subjected to Sar1 knockdowns do not fully develop, we opted to initially induce the knockdown using a mosaic Gal4 line (*otu*-Gal4). This Gal4 permits mild expression of Sar1, resulting in a less severe phenotype. By visualizing Me31B, Tral, and Cup in this background, we determined that only a limited number of P-bodies were present (Fig. 4A). This observation was surprising since a study in yeast found that in the absence of Sar1, there was an increase in P-body number (28). This difference may serve as evidence of the various pathways of P-body formation that are implemented across species and cell types.

**Figure 4:**
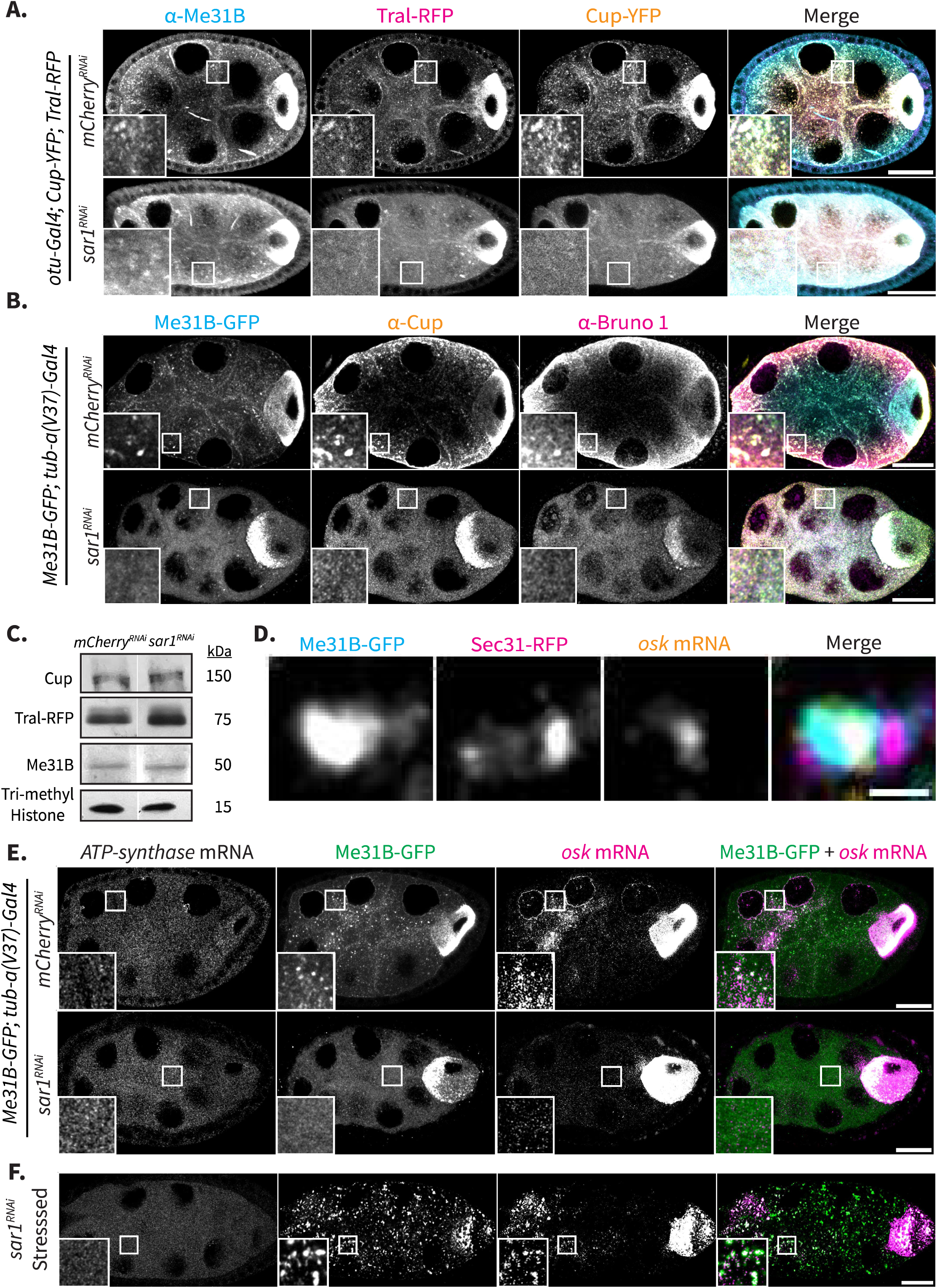
In the absence of ER exit sites, P-body integrity is compromised. **(A)** Co-visualization of Me31B, Tral-RFP, and Cup-YFP in *mCherry^RNAi^* and *sar1^RNAi^* egg chambers. **(B)** Me31B-GFP co-visualized with Cup and Bruno 1 in *mCherry^RNAi^* and *sar1^RNAi^* egg chambers. Scale bars are 20*µ*m. **(C)** Western blot analysis of Cup, Tral-RFP, and Me31B (Tri-methyl-Histone -- loading control). **(D)** Me31B-GFP condensate colocalized with Sec31-RFP and *oskar* mRNA visualized with smFISH probes. Scale bar is 1*µ*m. **(E)** Me31B-GFP, *ATP synthase* mRNA, and *oskar* mRNA visualized with smFISH probes in *mCherry^RNAi^* and *sar1^RNAi^* egg chambers. **(F)** Me31B-GFP, *ATP synthase* mRNA, and *oskar* mRNA visualized under nutritional stress in *sar1^RNAi^* background. All images are XY projections of 5 optical Z slices of 0.3*µ*m. Scale bars are 20*µ*m.

To determine if the decrease in Sar1 was responsible for the limited number of P-bodies, we visualized putative P-body proteins: Me31B, Cup, and Bruno 1, in the background of a strong Gal4 line (*a-tub-*Gal4) that induced a more substantial knockdown of Sar1. We observed that each protein was completely diffused in the nurse cell cytoplasm (Fig. 4B). Due to constraints of the *D. melanogaster* genome, we were unable to visualize Tral-RFP in this background. Notably, the stronger Gal4 line resulted in the complete diffusion of P-body components, while the weaker Gal4 line led to only an attenuation of P-body number. This finding is noteworthy, as it establishes a robust, causative link between ERES formation and P-body formation, which had not been previously demonstrated.

To determine if the reduced P-body population was a consequence of lower P-body protein concentrations in the *sar1^RNAi^* egg chambers, we conducted a Western blot analysis (Fig. 4C). Interestingly, P-body protein levels did not significantly change, suggesting that the lack of P-bodies in the Sar1 knockdown is not attributable to alterations in P-body protein levels, but rather to modifications in cytoplasmic protein distributions.

As many maternal mRNA’s are recruited into P-bodies, we further assessed whether their distribution was affected by the ERES knockdown. First, we aimed to establish the presence of putative P-body mRNAs within ERES P-bodies, as that may imply a reliance on ERES for mRNA granule formation. To investigate this, we co-visualized Me31B, Sec31, and several maternal mRNAs. Significantly, our findings revealed the colocalization of *oskar, nanos,* and *bicoid* mRNAs with ERES P-bodies (Fig. 4D, S4B, and S4C). We next visualized *ATP synthase* mRNA, a housekeeping transcript that does not associate with P-bodies, and *oskar* mRNA, a known P-body member, in *sar1^RNAi^* egg chambers (42,50). Notably, we found that the expression level and distribution of *ATP synthase* mRNA was unaffected, while *oskar* mRNA no longer formed large puncta in the nurse cell cytoplasm and was mislocalized to the anterior of the oocyte. This demonstrates that ERES are essential for the assembly of *oskar* mRNA into granules and, consequently, for its proper localization within the oocyte (Fig. 4E).

Given the shared characteristics between P-bodies and stress granules (i.e. composed of similar proteins and mRNAs, form via LLPS, and are expressed in the cytoplasm), we wanted to ascertain if disruption of ERES similarly impacted stress granule formation (51). To investigate this, we subjected females to starvation to induce stress, and visualized Me31B, *ATP synthase* mRNA, and *oskar* mRNA in *sar1^RNAi^* egg chambers. Surprisingly, we observed that stress granules could still form in these egg chambers, suggesting that ERES are exclusively crucial for P-body formation (Fig. 4F).

### ER stress is communicated via P-bodies and not stress granules

Cells communicate various types of stress to a diverse array of biological condensates. Many stressors, including starvation and virginity, induce stress granule formation and slowing of translation (52,53). ER stress represents a distinct form of stress, but similarly results in phosphorylation of EIF2*ɑ* and consequently a decrease in translation (54). ER stress arises predominantly from the presence of misfolded proteins and triggers the Unfolded Protein Response (UPR), which in turn signals multiple pathways. This leads to an increase in ER chaperone levels, a decrease in the ER’s synthetic load (55), induction of the antioxidant response pathway (56), and targeted mRNA degradation at the ER (57). ER stress is primarily cytoprotective, however, if ER stress persists, it can trigger apoptosis, making the ER a key player in cellular decision making (58). Given the intricate connections between ER stress and various cellular pathways, we wondered if ER stress is also communicated to P-bodies via ERES.

To differentiate between stress granules and P-bodies, we utilized Rin and eIF4G, two key stress granule markers. We induced ER stress by treating ovaries with 5mM DTT for 30 min, which prevents disulfide bond formation resulting in misfolded proteins (59). Interestingly, we observed that under ER stress, P-bodies were 71% larger (0.259*µ*m^3^ compared to 0.151*µ*m^3^) and 16% more spherical when compared to control (Fig. 5A, 5B, and 5C). This observation is in agreement with previous studies conducted in *D. melanogaster* embryonic S2 cells which showed that ER stress induced by Thapsigargin, an inhibitor of ER Ca^2+^ ATPases which triggers UPR, resulted in larger P-bodies (60). Surprisingly, ER stress did not induce the formation of stress granules, suggesting that the ER signals stress via P-bodies.

**Figure 5:**
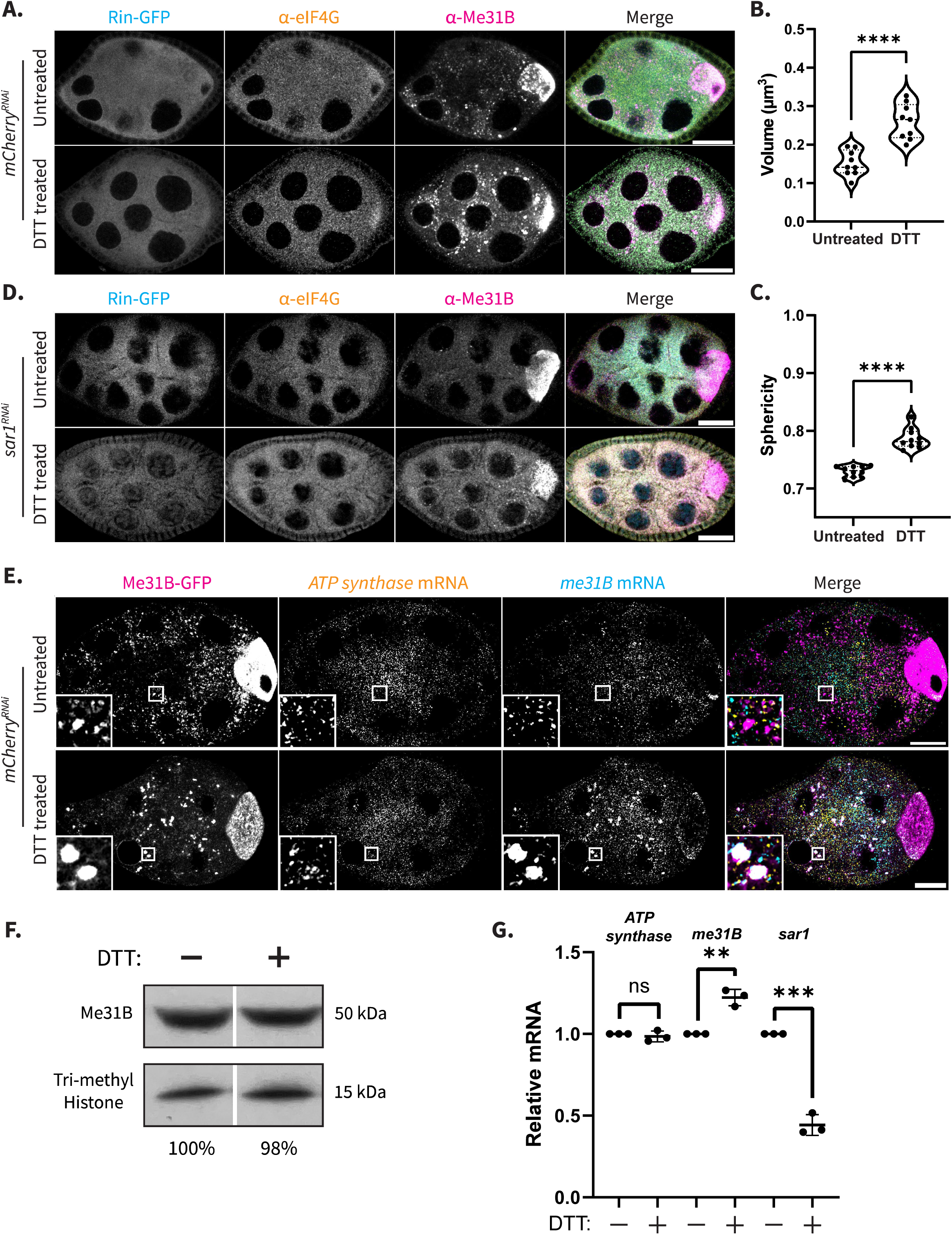
ER stress is communicated to the cell through P-bodies and not stress granules. **(A)** Co-visualization of Rin-GFP, eIF4G, and Me31B. ER stress induced in egg chambers with 30 min treatment with 5mM DTT. **(B,C)** Me31B condensate volume and sphericity measurements (n= 9). Significance calculated with Mann-Whitney statistical test. **(D)** Co-visualization of Rin-GFP, eIF4G, and Me31B in *sar1^RNAi^* egg chambers with induced ER stress. **(E)** Co-visualization of Me31B-GFP with *ATP synthase* and *me31B* mRNAs labeled with smFISH probes. **(F)** Western blot analysis of Me31B protein levels after 30 min DTT treatment compared to control. **(G)** RT-qPCR of *ATP synthase, me31B*, and *sar1* mRNAs in control and DTT treated egg chambers. Significance calculated with a t-test. **** p< .0001. All images are XY projections of 5 optical Z slices of 0.3*µ*m. Scale bars are 20*µ*m.

The absence of stress granule formation under ER stress was unexpected, particularly considering the readiness with which stress granules form in the *D. melanogaster* germline. To discern whether the absence of stress granules was due to ER stress directly or if the DTT treatment antagonizes stress granule integrity, we starved flies to induce stress granules and then subjected the ovaries to DTT treatment. We found that DTT did not lead to the dissolution of pre-formed stress granules, which suggests that DTT alone does not breakdown stress granules (Fig. S5).

To determine if ER stress was communicated to P-bodies through ERES specifically, we stressed *sar1^RNAi^* egg chambers with DTT. Interestingly, we found that, without ERES, Me31B-GFP still did not form granules, showing that it is via this interface that ER stress is communicated to membrane-less organelles (Fig. 5D).

To begin to elucidate the function of these ER stress-induced granules, we visualized *ATP synthase* and *me31B* mRNAs in DTT-treated egg chambers. *ATP synthase* mRNA does not enter stress granules or P-bodies as it is needed for normal cellular function. *me31B* mRNA similarly does not accumulate in P-bodies (61). Interestingly, under ER stress, both transcripts began to associate with the Me31B-GFP granules (Fig. 5E).

Transcripts recruited to granules can be translated at the periphery, stored, or degraded (62). Notably, the overall levels of Me31B protein were not affected by ER stress, suggesting that translation at the granules was not occurring (Fig. 5F). RT-qPCR revealed that *ATP synthase* mRNA levels stayed the same and *me31B* mRNA levels increased by ∼20%. This suggests that these Me31B-GFP granules are not sites of decay during ER stress, but perhaps function in mRNA storage (Fig. 5G). Interestingly, *sar1* mRNA levels decreased ∼50% after ER stress, which is consistent with ER stress leading to selective downregulation of genes involved in ER load (Fig. 5G). It is conceivable that these Me31B-GFP granules function in protecting essential transcripts during ER stress, potentially performing a more selective role than stress granules. However, further experiments would be necessary to assess this theory. Overall, this data indicates that ER stress alters P-body morphology, composition, and function in selective transcript storage.

### ERES knockdown leads to attenuation of P-body function

As P-bodies host multiple mRNA processes, we wanted to ascertain if these processes were compromised in the absence of ERES. We focused our studies on *oskar* mRNA as it forms a well-studied mRNP and is known to localize into P-bodies (42,63). First, we investigated ERES’s effect on translational repression by expressing endogenous Oskar-GFP in a *sar1^RNAi^* background. *oskar* mRNA is transcribed in the nurse cells throughout oogenesis and localizes posteriorly in the oocyte where Oskar protein is expressed during mid and late oogenesis. Interestingly, we found that, in *sar1^RNAi^* egg chambers, Oskar was prematurely expressed in early egg chambers, and only detectable within the oocyte (Fig. 6A).

**Figure 6:**
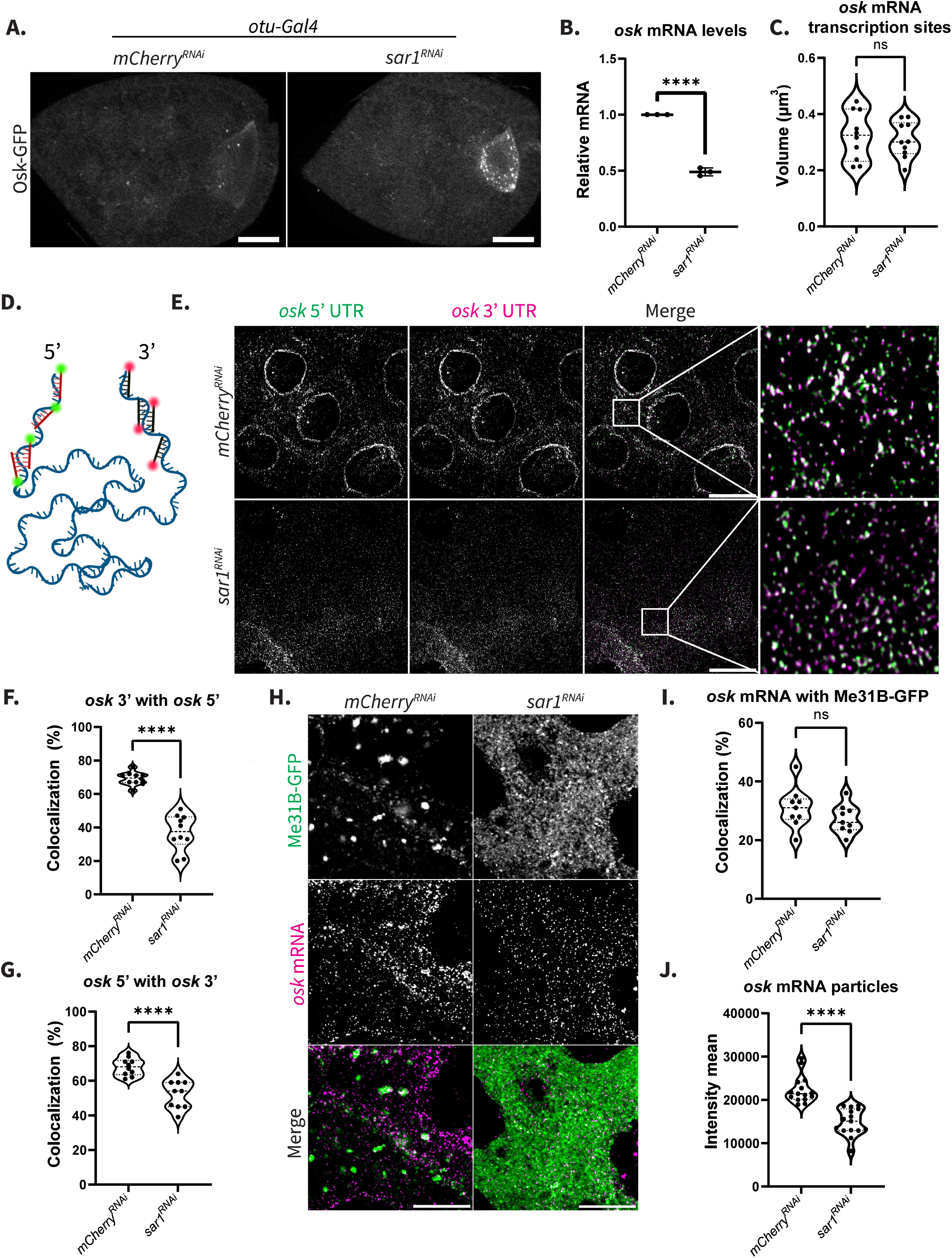
ERES knockdown leads to attenuation of P-body function. **(A)** Oskar-GFP visualized in *mCherry^RNAi^* and *sar1^RNAi^* egg chambers. XY projections of 5 optical Z slices of 0.3*µ*m. Scale bars are 20*µ*m. **(B)** *oskar* mRNA levels calculated with RT-qPCR in *mCherry^RNAi^* and *sar1^RNAi^*backgrounds. Significance calculated with a t-test. **(C)** Volume calculation of active transcription sites in *mCherry^RNAi^* and *sar1^RNAi^*egg chambers (n=10). **(D)** Diagram of 5’ UTR and 3’UTR *oskar* smFISH probes created with BioRender. **(E)** Co-visualization of *oskar* 5’ UTR and *oskar* 3’ UTR regions in *mCherry^RNAi^* and *sar1^RNAi^* egg chambers. XY projections of 5 optical Z slices of 0.3*µ*m. Scale bars are 20*µ*m. **(F,G)** Colocalization analysis of *oskar* 5’ UTR and *oskar* 3’ UTR (n=10). **(H)** Co-visualization of Me31B-GFP and *oskar* mRNA via STED in *mCherry^RNAi^* and *sar1^RNAi^* egg chambers. XY projections of 3 optical Z slices of 0.22*µ*m. Scale bars are 2*µ*m. **(I)** Colocalization analysis of *oskar* mRNA with Me31B-GFP (n= 9). **(J)** Intensity measurements of *oskar* mRNA particles in *mCherry^RNAi^* and *sar1^RNAi^* egg chambers (n= 15). Significance calculated with Mann-Whitney statistical test. **** p< .0001.

Next, we investigated the effect ERES have on P-body’s role in maintaining *oskar* mRNA stability. Using RT-qPCR, we found an average 51% decrease in *oskar* mRNA levels (Fig. 6B). To determine if this decrease was due to diminished RNA transcription or stability, we assessed the volume of active transcription sites in *sar1^RNAi^* egg chambers and determined that they were not significantly altered (Fig. 6C). To assess transcript stability, we used two sets of smFISH probes. One set detected the 5’ UTR of the *oskar* transcript and the other detected the 3’ UTR (Fig. 6D). Through imaging differentially-labeled *oskar* 5’ UTR and 3’ UTRs, followed by colocalization analysis, we observed that ∼40% more *oskar* transcripts were degraded in the *sar1^RNAi^*egg chambers compared to control. This suggests that the Sar1 knockdown did not result in decreased transcription in the nucleus, but rather in reduced transcript stability in the cytoplasm (Fig. 6E).

Using this method, we also established that *oskar* mRNA was degraded in both directions, but primarily underwent degradation via the 3’ to 5’ decay pathway (Fig. 6F and 6G). We performed this analysis for each of the COPII knockdowns and found a similar decrease in transcript stability, although not as pronounced as in Sar1 knockdown egg chambers (Fig. S6A-C). These findings strongly suggest that P-bodies play a significant role in mediating translational repression efficiency and transcript stability and these functions are compromised in the absence of ERES.

Given the established roles of P-body proteins Me31B, Cup, and Tral in translational repression and transcript stability, we asked whether compromised access of any of these proteins to the transcript may be responsible for the observed phenotypes (46,64). To address this possibility, we used STED super-resolution imaging to visualize Me31B-GFP, a key P-body protein in part responsible for *oskar* mRNA translational repression (63), together with *oskar* mRNA in *sar1^RNAi^* egg chambers. We found that the colocalization between *oskar* transcripts and Me31B-GFP was maintained (Fig. 6H and 6I). Cup similarly maintained association with *oskar* transcripts in *sar1^RNAi^* egg chambers (Fig. S6D and S6E). This indicates that it is the compromised P-body expression that contributes to an ease in translational repression and stability, rather than a breakdown at the mRNP level.

In support of this conclusion, we calculated the fluorescence intensity of *oskar* mRNA particles and found that in *sar1^RNAi^* egg chambers, *oskar* mRNA particles were less intense than in *mCherry^RNAi^* background. This indicates that *oskar* mRNA exists in lower-copy number mRNPs in the *sar1^RNAi^* background, compared to the *mCherry^RNAi^* background, where *oskar* mRNA is present in higher-copy number within these granules (Fig. 6J). This difference likely accounts for the observed decrease in the efficiency of translational repression and stability of *oskar* mRNA. Taken together this data suggests that translational repression and transcript stability are compromised in the absence of ERES due to attenuated P-body formation.

## DISCUSSION

Biological condensates are emerging as key regulators of cellular processes (9,34). While our understanding of their functions is expanding, a comprehensive picture of how membrane-bound and membrane-less organelles communicate remains largely unexplored. Here, we propose a mechanism wherein P-bodies and ERES work synergically to orchestrate spatial-temporal mRNA and protein localization. Previous studies have demonstrated the regulation of ERES localization in *D. melanogaster* by Tral (23). Here, we show the reciprocal regulation: ERES are necessary for proper P-body integrity during *D. melanogaster* oogenesis. We propose a model where ERES formation contributes to the organization of larger, gel-like P-bodies that serve as nucleating hubs from which the cytoskeletal network can pull away small, liquid-like mobile P-bodies (Fig. 7). Further biochemical studies would validate the link described in this proposed model, and we find the connection between an organelle responsible for significant cellular decision-making and P-bodies a compelling justification for further studies. Moreover, we find the prospect of founder *sar1* mRNPs that localize to the ER and dictate ERES localization to be an intriguing focus for investigation.

**Figure 7:**
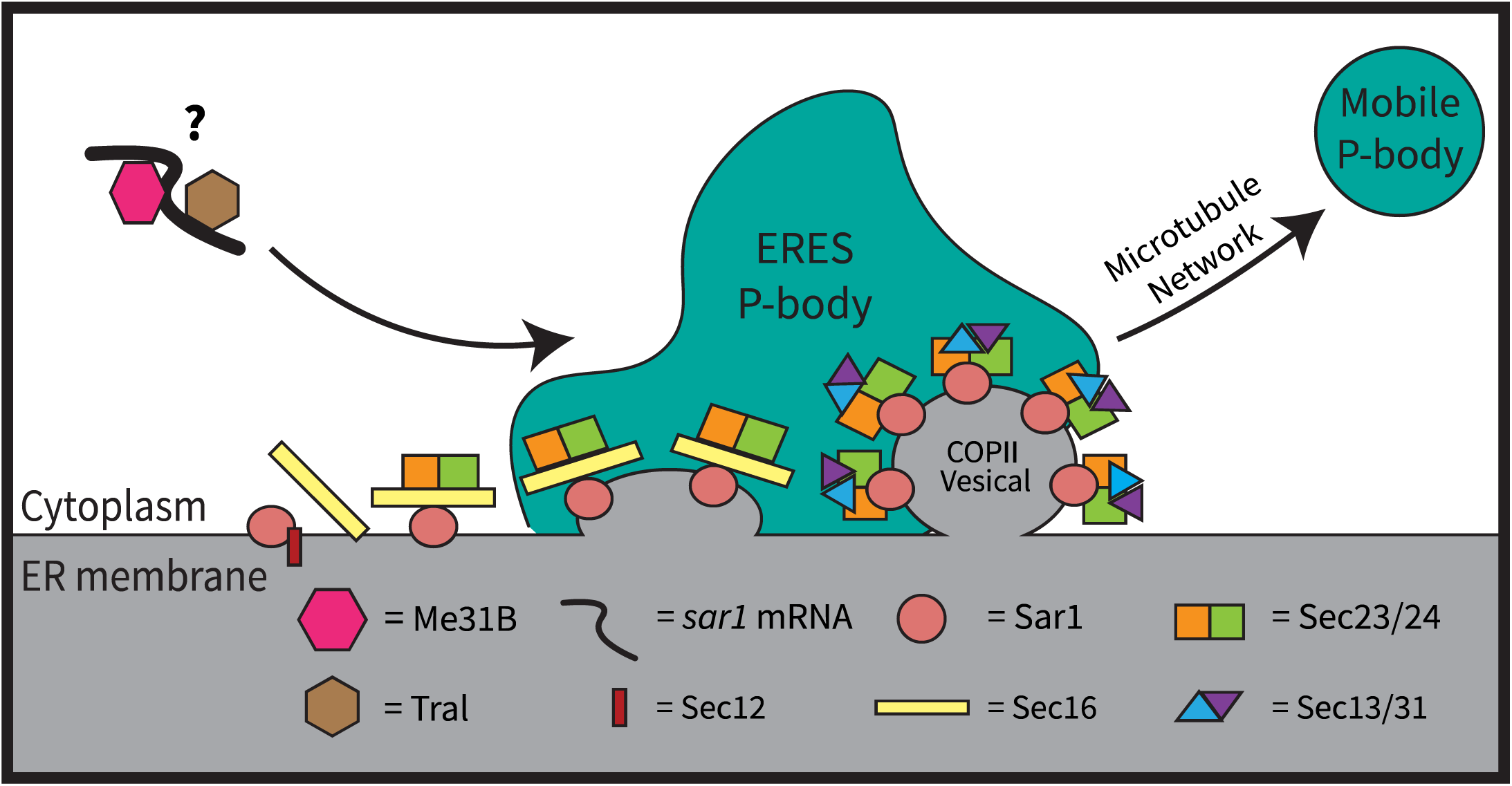
Proposed mechanism of P-body formation at ERES. ERES acting as a hub for P-body formation and maintenance. The model depicts a mature P-body docked at an ERES. Mobile, liquid-like P-bodies are pulled away from ERES hubs by the microtubule network. Founder *sar1* mRNA RNPs may be responsible for dictating localization of ERES P-body hubs.

### ERES P-bodies are a discrete class of P-bodies

Previous work has depicted P-bodies as a heterogeneous group of condensates, encompassing both liquid-like, mobile condensates, as well as solid-like, stationary condensates (65,66). Here, we demonstrate that this diverse population may actually represent distinct types of P-bodies with varying characteristics, potentially indicative of different functions. We propose that ERES P-bodies constitute one of these distinct classes.

Notably, the ER serves as a nucleating site for several LLPS bodies. TIS granules, membrane-less bodies that house a specific subset of mRNAs, form in contact with the ER membrane (67). Similarly, the assembly of Whi3 condensate in fungi is regulated, at least in part, by ER contact sites (68). Another type of LLPS condensate called Sec bodies also form at the ER under stress and are composed of ERES proteins (69). Since the presence of LLPS initiation at the ER is well established, we believe that the connection between P-body formation and ERES is a natural one. This idea is particularly compelling since ERES are involved in regulating long-distance protein localization, while P-bodies potentially work to regulate long-distance mRNA localization, making their functions somewhat parallel. The reciprocal regulation between P-bodies and ERES may serve a teleological function in establishing polarity over long distances.

### COPII vesicle components dictate P-body organization

P-bodies encompass a diverse array of proteins and transcripts. Me31B, Tral, and Cup are canonical P-body proteins that interact with each other and are interchangeably used as P-body markers. However, our findings reveal that these proteins are not uniformly affected by COPII knockdowns. While Me31B and Tral condensates exhibited similar alterations, Cup remained unaffected. This underscores the necessity for a comprehensive understanding of how each of the proteins present in P-bodies contributes to overall condensate state and function.

A notable observation was the nuanced roles played by each COPII protein in condensate organization. Sec16 appeared crucial for overall mature condensate formation, as evidenced by the smaller, more spherical Me31B and Tral condensates observed in its absence. The knockdown of coat proteins led to P-bodies that were more spherical, indicating a lack of proper, mature condensate formation. They were also larger, suggesting that they were improperly coalescing or undergoing coarsening processes like Ostwald ripening (70). Investigating the viscoelastic properties of these granules is essential to further discern the cause of the observed changes in morphology and size.

### P-bodies are necessary for optimal *oskar* mRNA regulation

Our findings suggest that P-bodies play a role in increasing the efficiency of translational repression and maintaining transcript stability. In the absence ERES P-body formation was compromised, and we observed increased *oskar* transcript degradation and improper translation. While there was a decrease in the efficiency of both pathways, translational repression and transcript stability were still mostly maintained in the absence of detectable P-bodies. This suggests that P-bodies are essential for optimal efficiency but are not indispensable to these pathways, which is in agreement with prior studies (71,72).

### ERES communicate with P-bodies

The secretory pathway plays a vital role in relaying external signals into the internal environment of the cell. An RNAi screen revealed that at least 122 kinases and phosphatases affect ER export. Among these are members of canonical EGF signaling pathways. The same study found that an increase in mitogen activated protein kinase (MAPK) pathways led to phosphorylation of Sec16 by ERK which increased ERES numbers (73). Since our data suggests that ERES influence P-body formation and maintenance, there may be an interesting connection between cancer and the interface between the ER and membrane-less organelles.

It is notable that our findings suggest that ER stress is communicated via P-bodies and not stress granules. ER stress is increasingly recognized as a significant promoter of pathology, underscoring the importance of elucidating the far-reaching effects of chronic ER stress. Our findings reveal that upon induction of ER stress, P-bodies form large, spherical condensates reminiscent of stress granules. This observation leads us to believe that these condensates are distinct from stress granules and are specifically triggered by ER stress. These granules accumulated both housekeeping mRNAs and mRNAs encoding a P-body protein, suggesting a potential role in altering post-transcriptional gene regulation during ER stress.

Here, we propose a paradigm where ERES serve as connection points between membrane-bound organelles and membrane-less P-bodies. What we find most compelling about this model is that it allows for positive feedback between P-bodies and ERES, offering an explanation for the abundance of both structures in some cell types compared to others. However, much remains to be understood about the relationship between these sites and biological condensates. Conducting studies in various tissues concurrently will be essential for gaining a comprehensive view of the far-reaching implications of this regulatory mechanism for cell viability.

## MATERIAL AND METHODS

### Fly husbandry

*Drosophila melanogaster* stocks were maintained on standard cornmeal agar food at 25°C. Female flies were put in grape vials and fed yeast paste 2-3 days prior to dissection. Fly stocks obtained from Bloomington *Drosophila* Stock Center: UAS-*mCherry^RNAi^* (BL #35785), *Me31B-GFP* (BL #52530), UAS*-KDEL-RFP* (BL #30910), UAS*-Sec31-RFP* (BL #86532), *tub-a*(V37)-Gal4 (BL #7063), UAS-*bicD^RNAi^* (BL #42929), UAS-*sec16^RNAi^* (BL #53917), UAS-*sec23^RNAi^* (BL #32365), UAS-*sec13^RNAi^* (BL #32468), UAS-*sec31^RNAi^* (BL #32878), UAS-*sar1^RNAi^* (BL #32364), *otu* -Gal4 (BL # 58424). Kyoto *Drosophila* Stock Center: *Cup-YFP* (DGRC 115-161). Vienna *Drosophila* Resource Center: *Rin-GFP* (VDCR #318907). *Oskar-GFP* stock was a generous gift from Dr. G. Gonsalves (Augusta University), and *Tral-RFP,* a generous gift from Dr. St. Johnston (Gurdon Institute at the University of Cambridge). UAS-*mCherry^RNAi^* (BL #35785) was used as a control in all RNAi experiments to account for effects of activated RNAi machinery. All RNAi lines were driven by *tub-a*(V37)-Gal4 (BL #7063) unless otherwise specified.

### Immunofluorescence

Ovaries were dissected and fixed in 2% PFA in PBS for 10 min. Egg chambers were washed 3X for 10 min in PBST (0.3% Triton X-100), permeabilized and blocked for 2 hrs in PBS with 1% Triton X-100 and 1% BSA and incubated with primary antibodies overnight at RT with rocking. Followed by 3X 10 min washes in PBST and incubation with fluorescently labeled secondary antibodies (1:1000; DyLight 488, 550, and 650; ThermoFisher). Followed by 3X 10 min washes in PBST and mounted in ProLong Diamond Antifade Mountant (Life Technologies). Antibodies used: Rabbit anti-Sec23 PA1-069A (1:200; ThermoFisher), Rabbit anti-BicD B11 deposited to the DSHB by Steward, R. (1:100; Developmental Studies Hybridoma Bank), Rabbit anti-eIF4G (1:200) kind gift from Dr. Izaurralde (Max Planck Institute for Developmental Biology), mouse anti-Cup (1:1000) and mouse anti-Me31B (1:1000) generous gifts from Dr. A. Nakamura (Institute of Molecular Embryology and Genetics, Kumamoto University). Rabbit-anti-Bruno 1 (1:8,000) kind gift from Dr. M. Lilly (NIH-National Institutes of Child Health and Development). Actin stain: Phalloidin Alexa Fluor 647 (1:200; Life Technologies). Nuclear membrane stain: wheat germ agglutinin CF405S (1: 200; Biotium).

### Tubulin staining and depolymerization

Ovaries were dissected into BRB80 (0.5M K-PIPES pH 6.8, 2M MgCl_2_, and 0.5M K-EGTA) and permeabilized in 1% Triton X-100 in BRB80 without rocking. They were rinsed in BRB80 and fixed with 2% PFA for 10 min. Egg chambers were washed 3X 10 min in PBST followed by 1X 10 min in 2X SSC 10% formamide, and incubated overnight in mouse anti-*ɑ*-Tubulin AA4.3-s deposited to the DSHB by Walsh, C. and anti-*ɑ*-Tubulin 12G10 deposited to the DSHB by Frankel, J. / Nelsen, E.M (1: 100; Developmental Studies Hybridoma Bank). Egg chambers were washed 3X 20 min in PBST and fixed in 4% PFA followed by 3X 10 min washes in PBST preceding a 2 hr incubation in a fluorescently labeled secondary (1:1000; Dylight 550 or 650; ThermoFisher). Washed 3X for 10 min in PBST and mounted as previously described. Microtubule depolymerization was achieved by teasing ovaries into Schneider’s media (ThermoFisher) supplemented with 10*µ*M colcemid (Enzo Life Sciences). Ovaries were rocked at RT for 30 min. Control ovaries were dissected into non-supplemented Schneider’s media.

### smFISH labeling

Ovaries were dissected and fixed in 4% PFA in PBS for 10 min. Egg chambers were washed 3X 10 min with 2X SSC and pre-hybridized with a 15 min wash in 2X SSC with 10% formamide before being incubated in smFISH probes (1:50) overnight at 37°C. Egg chambers were washed in pre-warmed 37°C 2X SSC with 10% formamide 3X 10 min and mounted as previously described. *oskar* 5’ UTR labeled with 46 probes and *oskar* 3’ UTR labeled with 45 probes.

### Microscopy

All imaging was performed on a Leica TCS SP8 Laser Scanning Microscope equipped with a white light laser (470-670nm), a solid-state laser (405nm), and a STED 660nm CW high intensity laser. Optical sections were acquired using an automated XYZ-piezo stage. All image files were saved as 16-bit data files. Acquisition software: Leica LAS-X.

### Live imaging

Ovaries were dissected into Halocarbon Oil 700 (Sigma-Aldrich) on a #1.5 coverslip. Egg chambers were adhered by pulling the tissue across the surface as described in (74).

For all live imaging, the 63X/1.4 oil objective was used. Each movie was acquired at 4X zoom in nurse cells in mid-stage egg chambers. Ten 0.3*µ*m optical Z sections were acquired every 15 seconds for 30 min and deconvolved using Leica’s Lightning module.

### Fixed imaging

Ovaries were dissected into 4% PFA in PBS and fixed for 10 min. They were washed 3X 10 min in PBST, 1X for 5 min in 0.1M glycine pH 3, and stored in 30% sucrose overnight 4°C. The following day they were flash frozen in O.C.T Compound (Tissue-Tek) before being sectioned on a cryotome.

For all laser scanning confocal imaging, the 63X/1.4 oil objective was used and optical Z slices were 0.3*µ*m. For all STED imaging, the 100X/1.4 oil objective was used, optical Z slices were 0.22*µ*m, and the zoom was 5X. STED samples were prepared from 25*µ*m ovary sections.

### Imaging analysis

Identical image acquisition was used for the control and each experimental slide. All images were deconvolved pre-processing using Leica’s Lightning module. Figure images were processed using Fiji/ImageJ (NIH) (75). Quantification was performed using Imaris Microscopy Image Analysis software (Oxford Instruments). P-bodies and ERES were detected using the Imaris ‘Surface’ module which allows for object-object statistics. *oskar* mRNA particles were detected with the ‘Spot’ module as 0.2*µ*m diameter spots. Colocalization was determined between ‘Spot-Spot’, ‘Surface-Surface’, and ‘Spot-Surface’ modules. For ‘Spot-Spot’ less than 0.2*µ*m distance between two spots was considered colocalization as the software measures from the middle of the spot and each spot had a 0.1*µ*m radius. For ‘Surface-Surface’ calculations a distance of less than 0*µ*m was considered colocalization as this module measures from surface edge to surface edge. For ‘Spot-Surface’ colocalization, a distance less than 0.1*µ*m was considered colocalization as the ‘Surface’ module measures from the edge of a surface to the center of a spot. For all analysis, images were taken from three slides each made from a separate fly cross. All images were processed using identical batch parameters and statistics were calculated using Prism 8 Software (GraphPad). All imaging statistics were assessed using Mann-Whitney statistical tests. Live imaging data was assessed using the Imaris’s tracking module. Tracking data was similarly assessed using Prism 8.

### Western blot analysis

Ten ovaries for each genotype were dissected directly into 95*µ*l of 2X Laemmli Sample Buffer (Bio-Rad) supplemented with 5*µ*l of BME and immediately mechanically lysed. They were heated at 95°C for 10 min and centrifuged for 10 min at 10,000 RCF. Lysates were loaded onto a 10% acrylamide gel. Antibodies used: mouse anti-Cup (1:3000), mouse anti-Me31B (1: 2000), Tral-RFP detected via Rabbit anti-GFP (1:10,000, Millipore), and Rabbit anti-Tri-methyl-Histone (C42D8) (1:150,000; Cell Signaling Technology). Bands visualized using TrueBlot ULTRA: Anti-Mouse and Anti-Rabbit Ig HRP (1:50,000; Rockland) with SuperSignal West Femto Maximum Sensitivity Substrate (ThermoFisher).

### RNA isolation and RT-qPCR

Whole ovaries were dissected into 4°C PBS. Ovaries were mechanically lysed in TRIzol (ThermoFisher) to extract total RNA. RNA was washed with ethanol and eluted in RNAse free water. Reverse Transcriptase reactions using 2.5*µ*g total RNA were performed using Superscript IV kit (Life Technologies). Primers were designed using DRSC FlyPrimerBank and made by Integrated DNA Technologies. RT-qPCR was performed with a Roche Lightcycler 480 (Roche Molecular Systems, Inc.). Each reaction contained 1*µ*l of cDNA, 4*µ*l of 10*µ*M primers, and 5*µ*l of SYBR Green 1 Master Mix (Roche Diagnostics, Indianapolis, IN). Statistical significance was determined via t-test.

### Stress Induction

For cellular stress induction via nutritional deprivation, flies were collected in grape vials and fed yeast paste for 2-3 days. 4 hrs prior to dissection, half the flies were moved to new grape vials with no yeast. Ovaries were then fixed and mounted as previously described. For ER stress induction, ovaries were teased into 5mM DTT (ThermoFisher) in Schneider’s media and incubated for 30 min. Control flies were incubated in Schneider’s media alone for 30 min. Both sets were then fixed and mounted as previously described.

## ACKNOWLEDGEMENTS

We thank Dr. A. Spralding (Carnegie Institution for Science), Dr. A. Nakamura, (Riken Center for Developmental Biology), Dr. M. Lilly (NIH), and Dr. E. Izaurralde (Max Planck Institute for Developmental Biology) for the kind gifts of antibodies. We extend thanks to Dr. D. St Johnston (University of Cambridge) and Dr. G. Gonsalves (Augustana University) for the kind gifts of *D. melanogaster* lines. We thank the BDSC Indiana and Kyoto DGRC for providing *D. melanogaster* lines, as well as the TRiP at Harvard Medical School (NIH/NIGMS RO1-GM084947) for the transgenic RNAi stocks. We thank the Bioimaging Facility at Hunter College for access to the Leica TCS SP8 and Imaris -- Image Analysis software. We thank Dr. P. Feinstein (Hunter College) for allowing us to use the Roche Light-cycler instrument.

## REFERENCES

1. Elbaum-Garfinkle S, Kim Y, Szczepaniak K, Chen CCH, Eckmann CR, Myong S, et al. The disordered P granule protein LAF-1 drives phase separation into droplets with tunable viscosity and dynamics. Proc Natl Acad Sci. 2015 Jun 9;112(23):7189–94.

2. Nott TJ, Petsalaki E, Farber P, Jervis D, Fussner E, Plochowietz A, et al. Phase Transition of a Disordered Nuage Protein Generates Environmentally Responsive Membraneless Organelles. Mol Cell. 2015 Mar;57(5):936–47.

3. Molliex A, Temirov J, Lee J, Coughlin M, Kanagaraj AP, Kim HJ, et al. Phase Separation by Low Complexity Domains Promotes Stress Granule Assembly and Drives Pathological Fibrillization. Cell. 2015 Sep;163(1):123–33.

4. Burke KA, Janke AM, Rhine CL, Fawzi NL. Residue-by-Residue View of In Vitro FUS Granules that Bind the C-Terminal Domain of RNA Polymerase II. Mol Cell. 2015 Oct;60(2):231–41.

5. Brangwynne CP, Eckmann CR, Courson DS, Rybarska A, Hoege C, Gharakhani J, et al. Germline P Granules Are Liquid Droplets That Localize by Controlled Dissolution/Condensation. Science. 2009 Jun 26;324(5935):1729–32.

6. Van Treeck B, Parker R. Emerging Roles for Intermolecular RNA-RNA Interactions in RNP Assemblies. Cell. 2018 Aug;174(4):791–802.

7. Schütz S, Nöldeke ER, Sprangers R. A synergistic network of interactions promotes the formation of in vitro processing bodies and protects mRNA against decapping. Nucleic Acids Res. 2017 Jun 20;45(11): 6911–22.

8. Borcherds W, Bremer A, Borgia MB, Mittag T. How do intrinsically disordered protein regions encode a driving force for liquid–liquid phase separation? Curr Opin Struct Biol. 2021 Apr;67:41–50.

9. Banani SF, Lee HO, Hyman AA, Rosen MK. Biomolecular condensates: organizers of cellular biochemistry. Nat Rev Mol Cell Biol. 2017 May;18(5):285–98.

10. Lin Y, Protter DSW, Rosen MK, Parker R. Formation and Maturation of Phase-Separated Liquid Droplets by RNA-Binding Proteins. Mol Cell. 2015 Oct;60(2):208–19.

11. Garaizar A, Espinosa JR, Joseph JA, Collepardo-Guevara R. Kinetic interplay between droplet maturation and coalescence modulates shape of aged protein condensates. Sci Rep. 2022 Mar 15;12(1):4390.

12. McLaughlin JM, Bratu DP. *Drosophila* melanogaster Oogenesis: An Overview. In: Bratu DP, McNeil GP, editors. Drosophila Oogenesis [Internet]. New York, NY: Springer New York; 2015 [cited 2024 Jun 11]. p. 1–20. (Methods in Molecular Biology; vol. 1328).

13. Lasko P. mRNA Localization and Translational Control in *Drosophila* Oogenesis. Cold Spring Harb Perspect Biol. 2012 Oct 1;4(10):a012294–a012294.

14. Sheth U, Parker R. Decapping and Decay of Messenger RNA Occur in Cytoplasmic Processing Bodies. Science. 2003 May 2;300(5620):805–8.

15. Standart N, Weil D. P-Bodies: Cytosolic Droplets for Coordinated mRNA Storage. Trends Genet. 2018 Aug 1;34(8):612–26.

16. Lin MD, Jiao X, Grima D, Newbury SF, Kiledjian M, Chou TB. *Drosophila* processing bodies in oogenesis. Dev Biol. 2008 Oct;322(2):276–88.

17. Zeitelhofer M, Macchi P, Dahm R. Perplexing bodies: The putative roles of P-bodies in neurons. RNA Biol. 2008 Oct;5(4):244–8.

18. Kurokawa K, Nakano A. The ER exit sites are specialized ER zones for the transport of cargo proteins from the ER to the Golgi apparatus. J Biochem (Tokyo). 2019 Feb 1;165(2):109–14.

19. Aridor M, Guzik AK, Bielli A, Fish KN. Endoplasmic Reticulum Export Site Formation and Function in Dendrites. J Neurosci. 2004 Apr 14;24(15):3770–6.

20. Aridor M, Fish KN. Selective Targeting of ER Exit Sites Supports Axon Development. Traffic. 2009 Nov;10(11):1669–84.

21. Herpers B, Rabouille C. mRNA Localization and ER-based Protein Sorting Mechanisms Dictate the Use of Transitional Endoplasmic Reticulum-Golgi Units Involved in Gurken Transport in *Drosophila* Oocytes. Mol Biol Cell. 2004 Dec;15(12):5306–17.

22. Decker CJ, Parker R. CAR-1 and Trailer hitch: driving mRNP granule function at the ER? J Cell Biol. 2006 Apr 24;173(2):159–63.

23. Wilhelm JE, Buszczak M, Sayles S. Efficient Protein Trafficking Requires Trailer Hitch, a Component of a Ribonucleoprotein Complex Localized to the ER in *Drosophila*. Dev Cell. 2005 Nov;9(5):675–85.

24. Kugler JM, Chicoine J, Lasko P. Bicaudal-C associates with a Trailer Hitch/Me31B complex and is required for efficient Gurken secretion. Dev Biol. 2009 Apr;328(1):160–72.

25. Squirrell JM, Eggers ZT, Luedke N, Saari B, Grimson A, Lyons GE, et al. CAR-1, a Protein That Localizes with the mRNA Decapping Component DCAP-1, Is Required for Cytokinesis and ER Organization in Caenorhabditis elegans Embryos. Mol Biol Cell. 2006;17.

26. Nguyen DTM, Koppers M, Farías GG. Endoplasmic reticulum – condensate interactions in protein synthesis and secretion. Curr Opin Cell Biol. 2024 Jun;88:102357.

27. Wu H, Carvalho P, Voeltz GK. Here, there, and everywhere: The importance of ER membrane contact sites. Science. 2018 Aug 3;361(6401):eaan5835.

28. Kilchert C, Weidner J, Prescianotto-Baschong C, Spang A. Defects in the Secretory Pathway and High Ca ^2+^ Induce Multiple P-bodies. Gilmore R, editor. Mol Biol Cell. 2010 Aug;21(15):2624–38.

29. Lee JE, Cathey PI, Wu H, Parker R, Voeltz GK. Endoplasmic reticulum contact sites regulate the dynamics of membraneless organelles. Science. 2020 Jan 31;367(6477):eaay7108.

30. Lee AS, Hendershot LM. ER stress and cancer. Cancer Biol Ther. 2006 Jul;5(7):721–2.

31. Lindholm D, Wootz H, Korhonen L. ER stress and neurodegenerative diseases. Cell Death Differ. 2006 Mar;13(3):385–92.

32. Luo Y, Na Z, Slavoff SA. P-Bodies: Composition, Properties, and Functions. Biochemistry. 2018 May;57(17):2424–31.

33. Villegas JA, Heidenreich M, Levy ED. Molecular and environmental determinants of biomolecular condensate formation. Nat Chem Biol. 2022 Dec;18(12):1319–29.

34. Lyon AS, Peeples WB, Rosen MK. A framework for understanding the functions of biomolecular condensates across scales. Nat Rev Mol Cell Biol. 2021 Mar;22(3):215–35.

35. Sankaranarayanan M, Emenecker RJ, Wilby EL, Jahnel M, Trussina IREA, Wayland M, et al. Adaptable P body physical states differentially regulate bicoid mRNA storage during early *Drosophila* development. Dev Cell. 2021 Oct;56(20):2886–2901.e6.

36. Shiina N. Liquid- and solid-like RNA granules form through specific scaffold proteins and combine into biphasic granules. J Biol Chem. 2019 Mar;294(10):3532–48.

37. Aizer A, Brody Y, Ler LW, Sonenberg N, Singer RH, Shav-Tal Y. The Dynamics of Mammalian P Body Transport, Assembly, and Disassembly In Vivo. Wickens M, editor. Mol Biol Cell. 2008 Oct;19(10):4154–66.

38. Baker FC, Neiswender H, Veeranan-Karmegam R, Gonsalvez GB. *In vivo* proximity biotin ligation identifies the interactome of Egalitarian, a Dynein cargo adaptor. Development. 2021 Nov 15;148(22):dev199935.

39. Van Der Verren SE, Zanetti G. The small GTPASE Sar1, control centre of COPII trafficking. FEBS Lett. 2023 Mar;597(6):865–82.

40. Hanna MG, Mela I, Wang L, Henderson RM, Chapman ER, Edwardson JM, et al. Sar1 GTPase Activity Is Regulated by Membrane Curvature. J Biol Chem. 2016 Jan;291(3):1014–27.

41. Ivan V, De Voer G, Xanthakis D, Spoorendonk KM, Kondylis V, Rabouille C. *Drosophila* Sec16 Mediates the Biogenesis of tER Sites Upstream of Sar1 through an Arginine-Rich Motif. Glick BS, editor. Mol Biol Cell. 2008 Oct;19(10):4352–65.

42. Bayer LV, Milano S, Formel SK, Kaur H, Ravichandran R, Cambeiro JA, et al. Cup is essential for *oskar* mRNA translational repression during early *Drosophila* oogenesis. RNA Biol. 2023 Dec 31;20(1):573–87.

43. Yoshihisa T, Barlowe C, Schekman R. Requirement for a GTPase-Activating Protein in Vesicle Budding from the Endoplasmic Reticulum. Science. 1993 Mar 5;259(5100):1466–8.

44. Bi X, Mancias JD, Goldberg J. Insights into COPII Coat Nucleation from the Structure of Sec23•Sar1 Complexed with the Active Fragment of Sec31. Dev Cell. 2007 Nov;13(5):635–45.

45. Antonny B, Madden D, Hamamoto S, Orci L, Schekman R. Dynamics of the COPII coat with GTP and stable analogues. Nat Cell Biol. 2001 Jun;3(6):531–7.

46. Nakamura A, Sato K, Hanyu-Nakamura K. *Drosophila* Cup Is an eIF4E Binding Protein that Associates with Bruno and Regulates *oskar* mRNA Translation in Oogenesis. Dev Cell. 2004 Jan;6(1):69–78.

47. Tritschler F, Eulalio A, Helms S, Schmidt S, Coles M, Weichenrieder O, et al. Similar Modes of Interaction Enable Trailer Hitch and EDC3 To Associate with DCP1 and Me31B in Distinct Protein Complexes. Mol Cell Biol. 2008 Nov;28(21):6695–708.

48. Tikhomirova MS, Kadosh A, Saukko-Paavola AJ, Shemesh T, Klemm RW. A role for endoplasmic reticulum dynamics in the cellular distribution of microtubules. Proc Natl Acad Sci. 2022 Apr 12;119(15):e2104309119.

49. Sweet TJ, Boyer B, Hu W, Baker KE, Coller J. Microtubule disruption stimulates P-body formation. RNA. 2007 Apr;13(4):493–502.

50. Kato Y, Nakamura A. Roles of cytoplasmic RNP granules in intracellular RNA localization and translational control in the *Drosophila* oocyte. Dev Growth Differ. 2012 Jan;54(1):19–31.

51. Corbet GA, Parker R. RNP Granule Formation: Lessons from P-Bodies and Stress Granules. Cold Spring Harb Symp Quant Biol. 2019;84:203–15.

52. Ivanov P, Kedersha N, Anderson P. Stress Granules and Processing Bodies in Translational Control. Cold Spring Harb Perspect Biol. 2019 May;11(5):a032813.

53. Protter DSW, Parker R. Principles and Properties of Stress Granules. Trends Cell Biol. 2016 Sep;26(9): 668–79.

54. Harding HP, Novoa I, Zhang Y, Zeng H, Wek R, Schapira M, et al. Regulated Translation Initiation Controls Stress-Induced Gene Expression in Mammalian Cells. Mol Cell. 2000 Nov;6(5):1099–108.

55. Schröder M, Kaufman RJ. ER stress and the unfolded protein response. Mutat Res Mol Mech Mutagen. 2005 Jan;569(1–2):29–63.

56. Cullinan SB, Diehl JA. PERK-dependent Activation of Nrf2 Contributes to Redox Homeostasis and Cell Survival following Endoplasmic Reticulum Stress. J Biol Chem. 2004 May;279(19):20108–17.

57. Hollien J, Weissman JS. Decay of Endoplasmic Reticulum-Localized mRNAs During the Unfolded Protein Response. Science. 2006 Jul 7;313(5783):104–7.

58. Gorman AM, Healy SJM, Jäger R, Samali A. Stress management at the ER: Regulators of ER stress-induced apoptosis. Pharmacol Ther. 2012 Jun;134(3):306–16.

59. Oslowski CM, Urano F. Measuring ER Stress and the Unfolded Protein Response Using Mammalian Tissue Culture System. In: Methods in Enzymology [Internet]. Elsevier; 2011 [cited 2024 Mar 4]. p. 71–92.

60. Loschi M, Leishman CC, Berardone N, Boccaccio GL. Dynein and kinesin regulate stress-granule and P-body dynamics. J Cell Sci. 2009 Nov 1;122(21):3973–82.

61. de Valoir T, Tucker MA, Belikoff EJ, Camp LA, Bolduc C, Beckingham K. A second maternally expressed *Drosophila* gene encodes a putative RNA helicase of the “DEAD box” family. Proc Natl Acad Sci. 1991 Mar 15;88(6):2113–7.

62. Weil TT, Parton RM, Herpers B, Soetaert J, Veenendaal T, Xanthakis D, et al. *Drosophila* patterning is established by differential association of mRNAs with P bodies. Nat Cell Biol. 2012 Dec;14(12):1305–13.

63. Nakamura A, Amikura R, Hanyu K, Kobayashi S. Me31B silences translation of oocyte-localizing RNAs through the formation of cytoplasmic RNP complex during *Drosophila* oogenesis. Development. 2001 Sep 1;128(17):3233–42.

64. Götze M, Dufourt J, Ihling C, Rammelt C, Pierson S, Sambrani N, et al. Translational repression of the Drosophila nanos mRNA involves the RNA helicase Belle and RNA coating by Me31B and Trailer hitch.

65. Aizer A, Shav-Tal Y. Intracellular trafficking and dynamics of P bodies. Prion. 2008 Oct;2(4):131–4.

66. Ditlev JA, Case LB, Rosen MK. Who’s In and Who’s Out—Compositional Control of Biomolecular Condensates. J Mol Biol. 2018 Nov;430(23):4666–84.

67. Ma W, Mayr C. A Membraneless Organelle Associated with the Endoplasmic Reticulum Enables 3*′*UTR-Mediated Protein-Protein Interactions. Cell. 2018 Nov;175(6):1492–1506.e19.

68. Snead WT, Jalihal AP, Gerbich TM, Seim I, Hu Z, Gladfelter AS. Membrane surfaces regulate assembly of ribonucleoprotein condensates. Nat Cell Biol. 2022 Apr;24(4):461–70.

69. Zacharogianni M, Aguilera-Gomez A, Veenendaal T, Smout J, Rabouille C. A stress assembly that confers cell viability by preserving ERES components during amino-acid starvation. eLife. 2014 Nov 11;3:e04132.

70. Berry J, Brangwynne CP, Haataja M. Physical principles of intracellular organization via active and passive phase transitions. Rep Prog Phys. 2018 Apr 1;81(4):046601.

71. Eulalio A, Behm-Ansmant I, Schweizer D, Izaurralde E. P-Body Formation Is a Consequence, Not the Cause, of RNA-Mediated Gene Silencing. Mol Cell Biol. 2007 Jun;27(11):3970–81.

72. Blake LA, Watkins L, Liu Y, Inoue T, Wu B. A rapid inducible RNA decay system reveals fast mRNA decay in P-bodies. Nat Commun. 2024 Mar 28;15(1):2720.

73. Farhan H, Wendeler MW, Mitrovic S, Fava E, Silberberg Y, Sharan R, et al. MAPK signaling to the early secretory pathway revealed by kinase/phosphatase functional screening. J Cell Biol. 2010 Jun 14;189(6):997–1011.

74. Prasad M, Wang X, He L, Cai D, Montell DJ. Border Cell Migration: A Model System for Live Imaging and Genetic Analysis of Collective Cell Movement. In: Bratu DP, McNeil GP, editors. Drosophila Oogenesis: Methods and Protocols [Internet]. New York, NY: Springer New York; 2015. p. 89–97.

75. Schindelin J, Arganda-Carreras I, Frise E, et al. Fiji: an open-source platform for biological-image analysis. Nat Methods. 2012;9(7):676–682.

