## Supplemental Figures for "The role of ER exit sites in maintaining P-body organization and transmitting ER stress response during *Drosophila melanogaster* oogenesis"

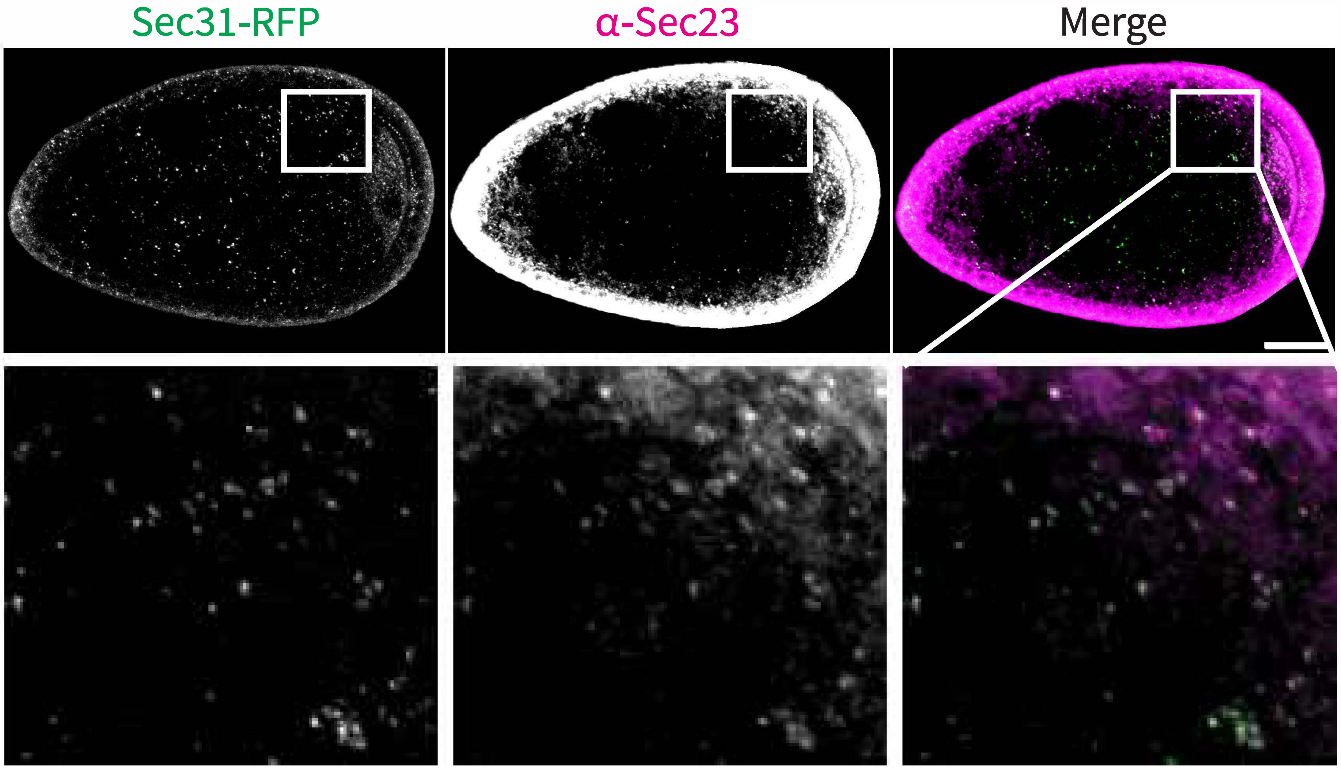
Sec31-RFP
α-Sec23
Merge

**Figure S1:** Co-visualization of Sec31-RFP with Sec23. XY projections of 5 optical Z slices of 0.3µm. Scale bars are 20µm and 3.3µm respectively in zoomed inset.

### Figure S2

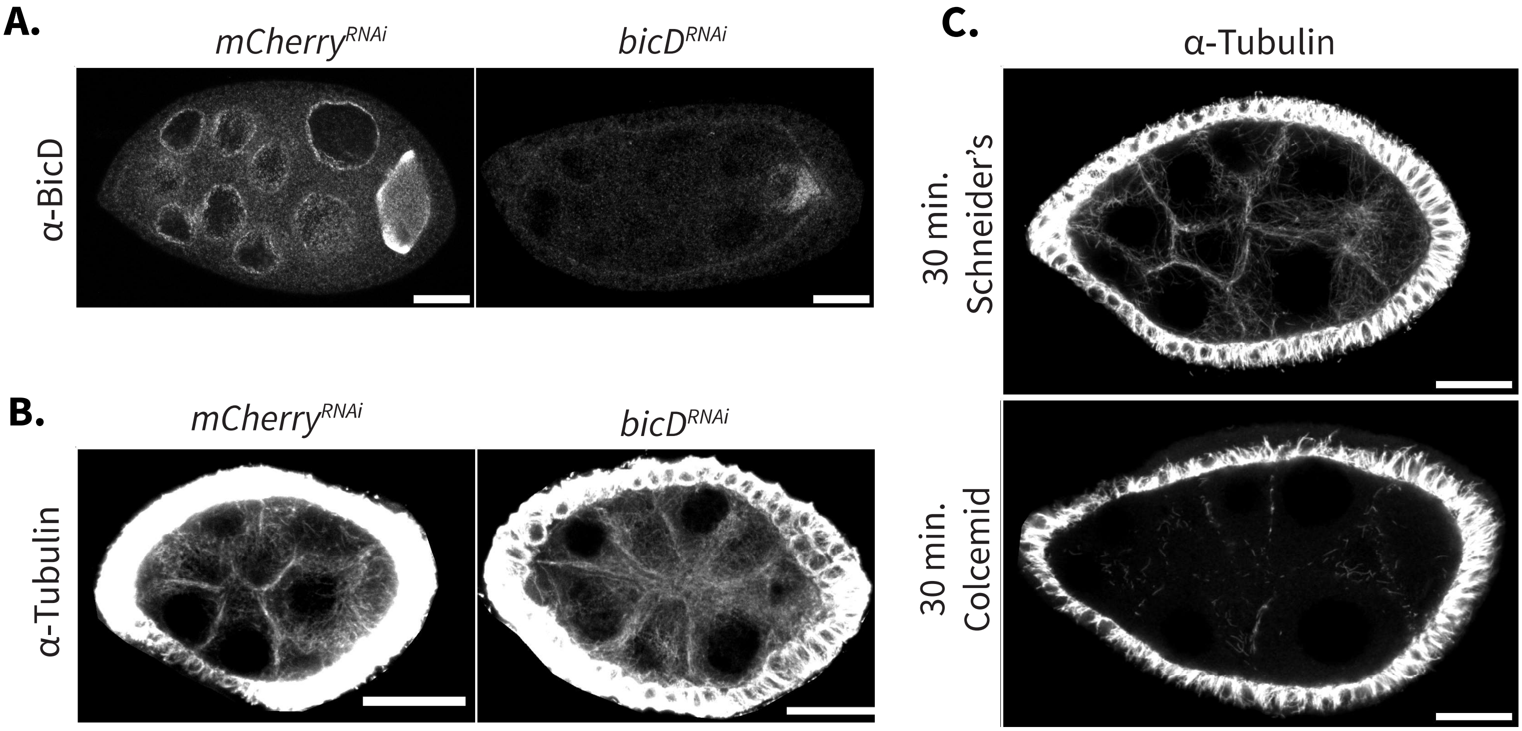
A.
mCherryRNAi
bicDRNAi
α-BicD
B.
mCherryRNAi
bicDRNAi
α-Tubulin
C.
α-Tubulin
30 min.
Schneider's
30 min.
Colcemid

**Figure S2:** **(A)** BicD expression in *mCherry*$^{RNAi}$ and *bicD*$^{RNAi}$ egg chambers. **(B)** α-Tubulin labeled microtubules in *mCherry*$^{RNAi}$ and *bicD*$^{RNAi}$ egg chambers. **(C)** Microtubule integrity in egg chambers after 30 mins incubation in Schneider's media or 10mM colcemid solution. All images are XY projections of 5 optical Z slices of 0.3μm. Scale bars are 20μm.

**Figure S3**



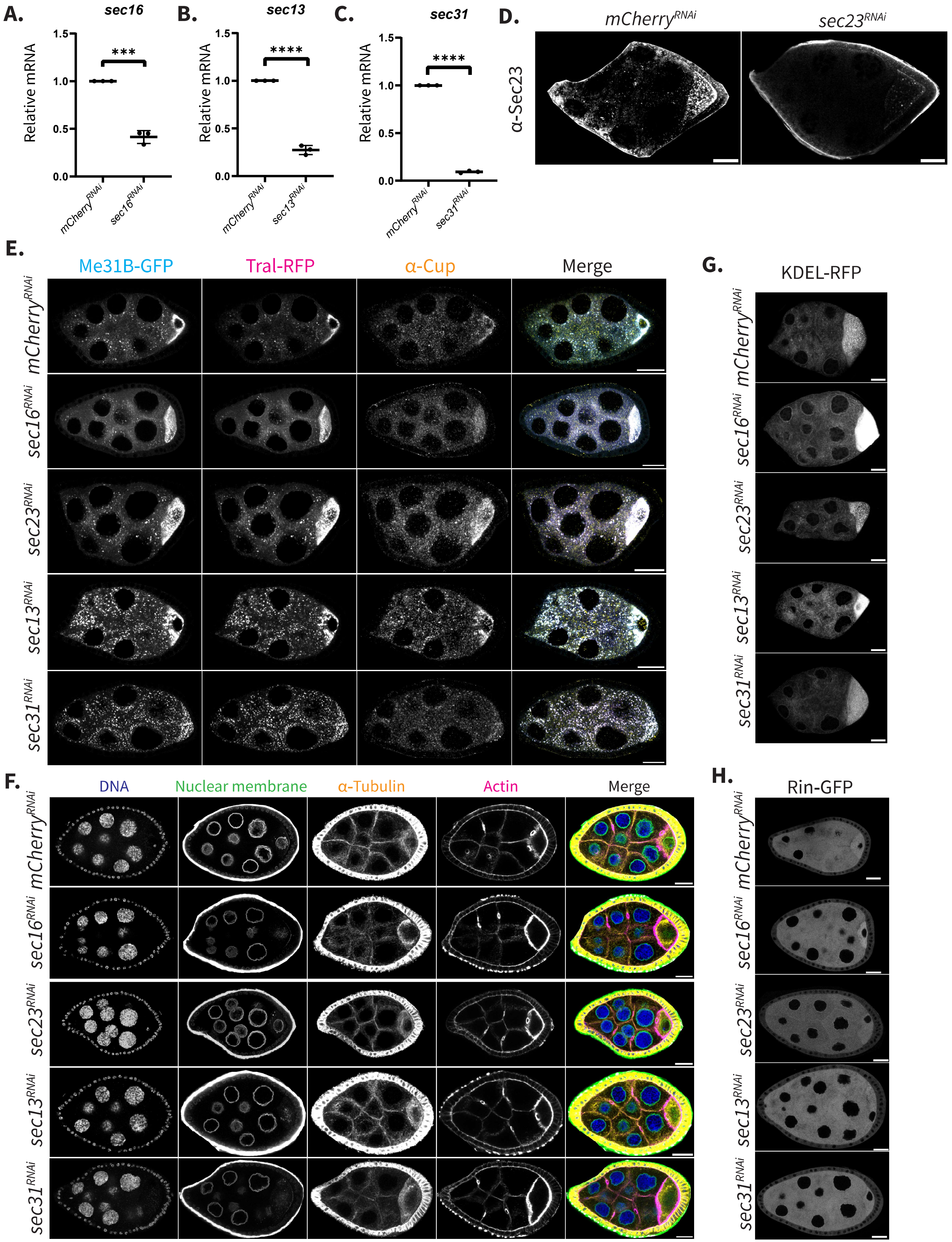
A.
sec16
Relative mRNA
1.5
1.0
0.5
0.0
***
mCherry^RNAi
sec16^RNAi
B.
sec13
Relative mRNA
****
mCherry^RNAi
sec13^RNAi
C.
sec31
Relative mRNA
****
mCherry^RNAi
sec31^RNAi
D.
mCherry^RNAi
sec23^RNAi
α-Sec23
E.
Me31B-GFP
Tral-RFP
α-Cup
Merge
mCherry^RNAi
sec16^RNAi
sec23^RNAi
sec13^RNAi
sec31^RNAi
F.
DNA
Nuclear membrane
α-Tubulin
Actin
Merge
G.
KDEL-RFP
H.
Rin-GFP

**Figure S3: (A,B,C)** Assessing knockdown efficiency of each ERES component via RT-qPCR. Significance calculated with a t-test. **** $p < .0001$. **(D)** Detection of Sec23 via antibody in *mCherry*$^{RNAi}$ and *sec23*$^{RNAi}$ egg chambers. **(E)** Full egg chamber images of Fig. 3B. **(F)** DNA (DAPI), nuclear membranes (wheat germ agglutinin), microtubules (α-Tubulin), and actin (phalloidin) in COPII component knockdown backgrounds. **(G)** KDEL-RFP visualized in each ERES component knockdown background. **(H)** Rin-GFP visualized in each ERES component knockdown background. All images are XY projections of 5 optical Z slices of 0.3µm. Scale bars are 20µm.

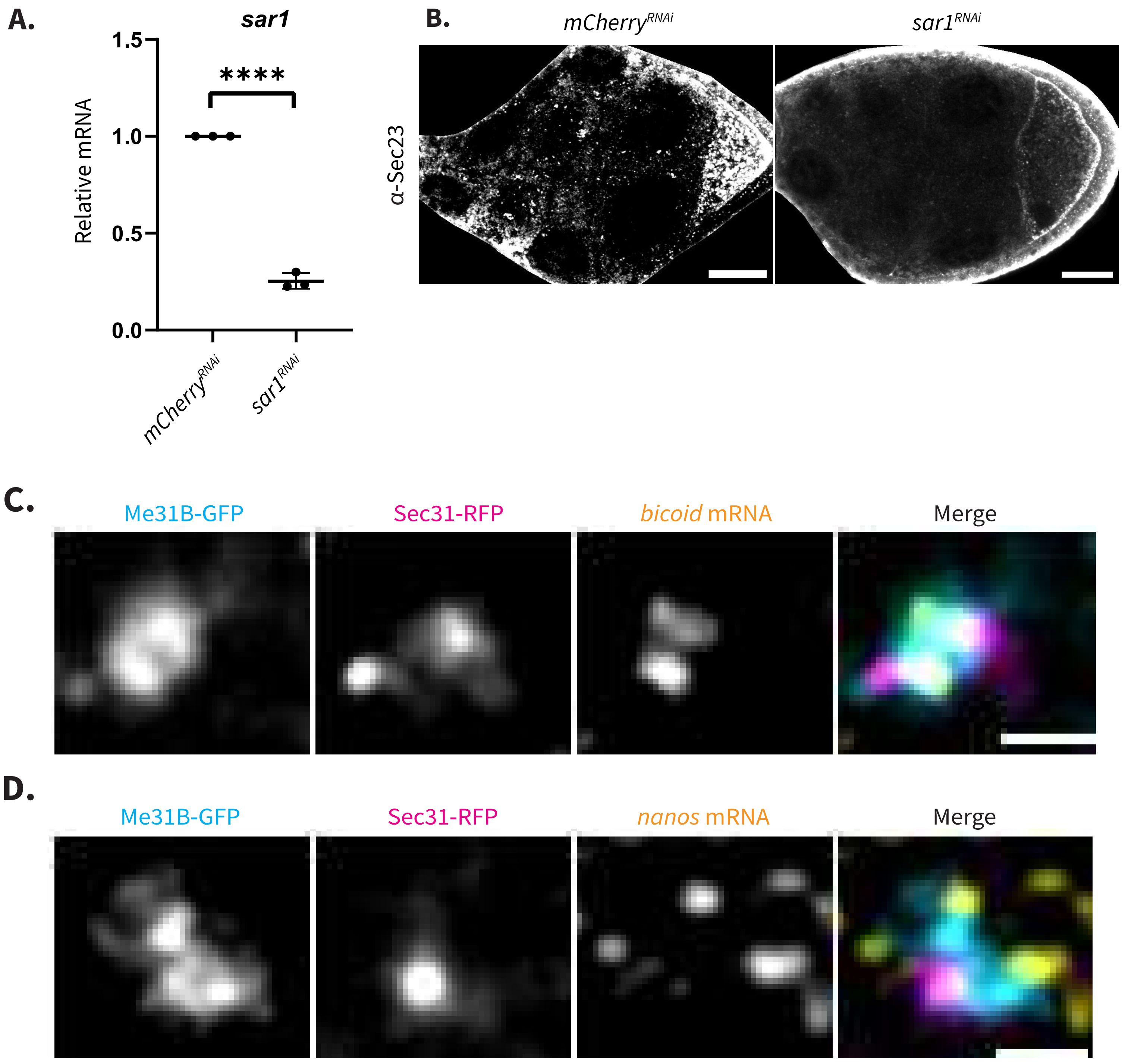
A.
sar1
1.5
1.0
0.5
0.0
Relative mRNA
****
mCherry^RNAi
sar1^RNAi
B.
mCherry^RNAi
sar1^RNAi
α-Sec23
C.
Me31B-GFP
Sec31-RFP
bicoid mRNA
Merge
D.
Me31B-GFP
Sec31-RFP
nanos mRNA
Merge

**Figure S4:** **(A)** Relative levels of *sar1* mRNA in *sar1*$^{RNAi}$ egg chamber detected with RT-qPCR. Significance calculated via a t-test. **** p< .0001 **(B)** Sec23 visualized in *mCherry*$^{RNAi}$ and *sar1*$^{RNAi}$ egg chambers. Scale bars are 20µm. **(C)** Me31B-GFP, Sec31-RFP, and *bicoid* mRNA visualized with smFISH probes. **(D)** Me31B-GFP, Sec31-RFP, and *nanos* mRNA visualized with smFISH probes. Scale bars are 1µm. All images are XY projections of 5 optical Z slices of 0.3µm.

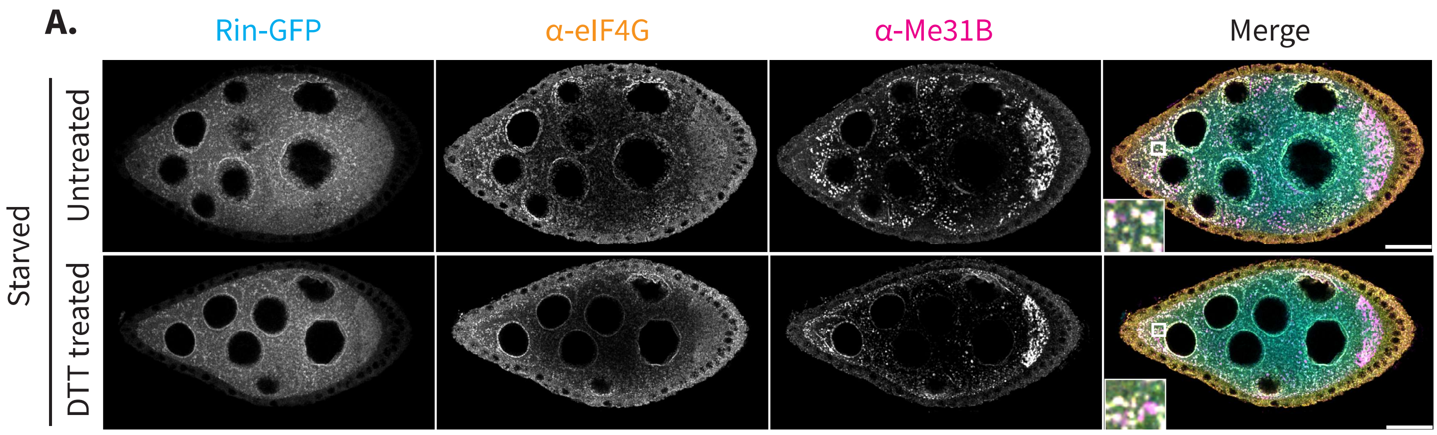
A.
Rin-GFP
α-eIF4G
α-Me31B
Merge
Starved
Untreated
DTT treated

**Figure S5:** Rin-GFP with eIF4G and Me31B visualized in egg chambers that were nutritionally stressed followed by induction of ER stress with DTT.

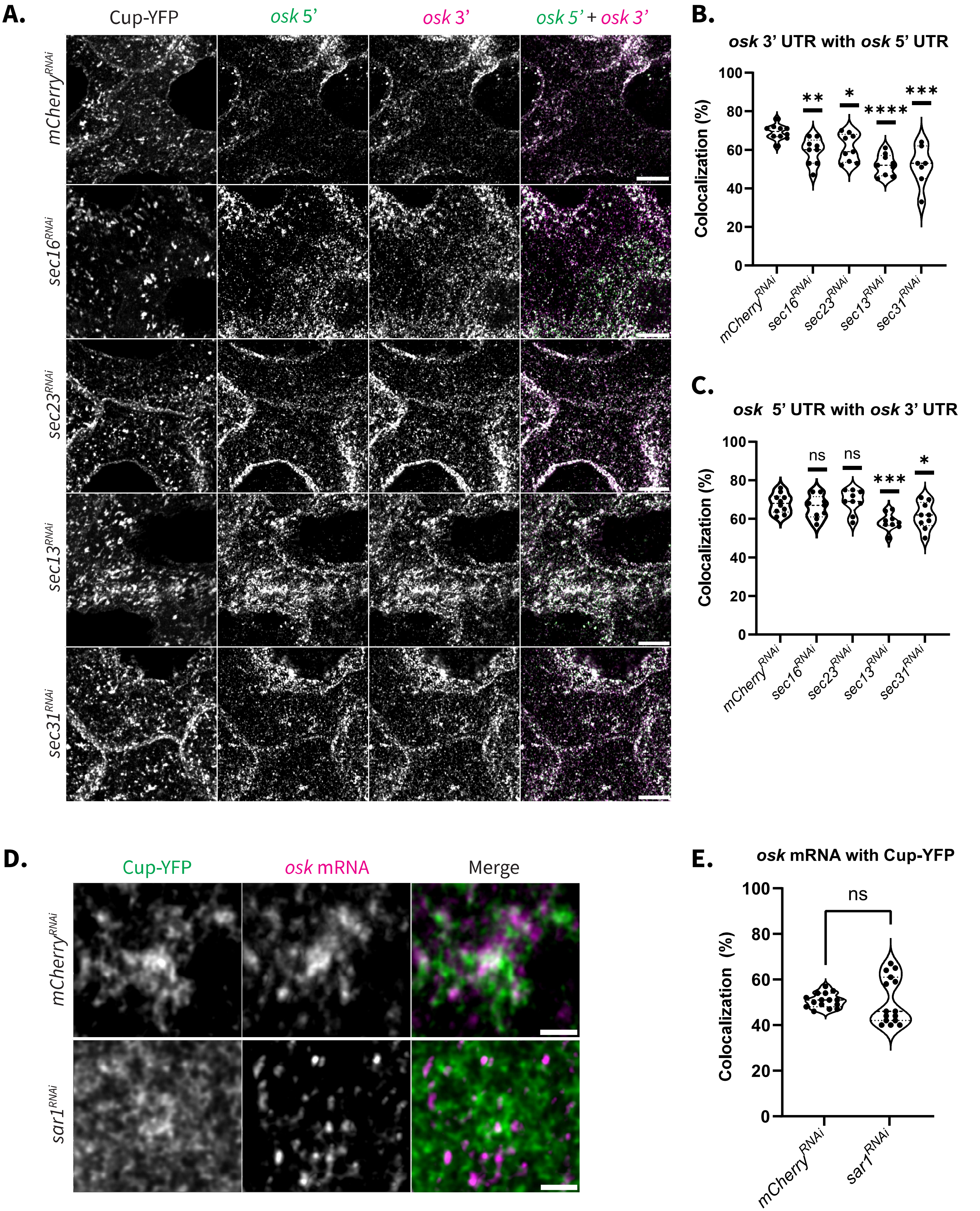
A.
Cup-YFP
osk 5'
osk 3'
osk 5' + osk 3'
mCherry RNAi
sec16 RNAi
sec23 RNAi
sec13 RNAi
sec31 RNAi
B.
osk 3' UTR with osk 5' UTR
Colocalization (%)
100
80
60
40
20
0
**
*
****
***
mCherry RNAi
sec16 RNAi
sec23 RNAi
sec13 RNAi
sec31 RNAi
C.
osk 5' UTR with osk 3' UTR
Colocalization (%)
100
80
60
40
20
0
ns
ns
***
*
mCherry RNAi
sec16 RNAi
sec23 RNAi
sec13 RNAi
sec31 RNAi
D.
Cup-YFP
osk mRNA
Merge
mCherry RNAi
sar1 RNAi
E.
osk mRNA with Cup-YFP
Colocalization (%)
100
80
60
40
20
0
ns
mCherry RNAi
sar1 RNAi

**Figure S6: (A)** Cup-YFP, *oskar* 5' UTR, and *oskar* 3' UTR visualized with smFISH probes in each of the COPII knockdown backgrounds. XY projections of 5 optical Z slices of 0.3µm. Scale bars are 20µm.
**(B,C)** Colocalization analysis of *oskar* 3' UTR and *oskar* 5' UTR, and colocalization analysis of *oskar* 5' UTR and *oskar* 3' UTR respectively (n=10). **(D)** STED acquired images of Cup-YFP and *oskar* mRNA.
XY projections of 5 optical Z slices of 0.22µm. Scale bars are 2µm. **(E)** Colocalization analysis of *oskar* mRNA with Cup-YFP (n= 15). Significance calculated with a Mann-Whitney statistical test. **** p< .0001.
